# Scalable Assembly of *Ascaris* Mitogenomes from Whole-Genome Data Reveals a Novel Clade

**DOI:** 10.64898/2025.12.18.695196

**Authors:** Lauren Woolfe, Kezia Kozel, Poom Adisakwattana, Allen Jethro Alonte, Kennesa Klariz Llanes, Alexandra Juhász, J. Russell Stothard, Christina Strube, Marie-Kristin Raulf, Scott P. Lawton, Toby Landeryou, Vachel Gay Paller, Umer Chaudhry, Arnoud H.M. van Vliet, Martha Betson

## Abstract

The genus *Ascaris* is an important group of giant parasitic roundworms, infecting over 700 million people globally, with substantial economic losses in domestic pigs. Whilst species of *Ascaris* are morphologically indistinguishable, analysis of mitochondrial and selected nuclear DNA loci has revealed three clades (A, B, C), which may inflate with the addition of genomic data. Here, we present a bioinformatic pipeline for *de novo* assembly of complete mitochondrial genomes (mtDNA) from low-coverage whole-genome data through host-read depletion or mtDNA read enrichment, followed by mtDNA-specific assembly.

Our approach yielded 149 high-quality *Ascaris* mtDNA assemblies, enabling the study of population-level diversity, including the identification of a novel clade (Clade D, designated here) associated with human samples from Ethiopia. Our analysis further revealed Clade C to comprise of pig-derived samples from Europe based on characterisation of worms isolated in Germany. Furthermore, genetic diversity across all mitochondrial loci was high (concatenated dataset: S = 1261; h = 94; π = 0.0224), indicating substantial intraspecies variation. Our methods described here provide a scalable framework for mtDNA genome reconstruction with insights into roundworm population-genomic and phylogenetic studies.

## 1. INTRODUCTION

Nematodes are a diverse group of worms capable of inhabiting a wide variety of ecological niches, including soil, fresh and saltwater ecosystems, and parasitise a broad range of plants and animals, including humans [1–3]. Parasitic nematodes are responsible for substantial morbidity and mortality in affected populations and have global economic implications, particularly in agriculture and healthcare [1,4–6]. Recognising their significant public health impact, the World Health Organisation has specifically targeted soil-transmitted helminths (STHs)—a subgroup of nematodes comprising genera such as *Ascaris*, *Trichuris*, *Ancylostoma* and *Necator* —for elimination as a public health problem by 2030 [7]. Concurrently, veterinary medicine faces increasing challenges from helminth infections, especially in livestock and companion animals, where the rise of anthelmintic resistance is resulting in reduced treatment efficacy, compromised animal welfare, decreased productivity, and economic losses [8–10].

Genomic markers provide powerful tools for detecting genetic variation, reconstructing evolutionary relationships, and monitoring phenotypic traits such as drug resistance or adaptation to environmental stressors [11–14]. In recent years, mitochondrial genomes (mitogenomes) have emerged as highly valuable genetic markers across diverse taxa due to their compact size, elevated copy number relative to nuclear DNA and conserved gene content [15,16].

Typically, nematode mitochondrial DNA (mtDNA) is composed of a single circular genome encoding 12 protein-coding genes, 2 ribosomal RNA (rRNA) genes, and 22 transfer RNA (tRNA) genes. While gene size and organisation are generally well conserved across nematodes [17], mtDNA evolves at rates up to ten times faster than nuclear DNA [16] resulting in significant variation driven by strong directional and multi-level selection linked to life cycles and environmental niches. The evolutionary patterns emerging from these selective processes are reflected in the phylogenetic relationships observed among nematode species. Owing to its lack of recombination and maternal [18–21] and high degree of genomic plasticity [17], mtDNA markers have become increasingly important in DNA barcoding (using standardized gene fragments, such as cytochrome oxidase subunit 1 [*cox1*] and NADH dehydrogenase subunit 1 [*nad1*], to distinguish between closely related lineages or genotypes), evolutionary studies, and species identification [2].

The most prevalent STH in humans is the giant roundworm *Ascaris,* which once matured can reach up to 30cm in length and resides in the small intestine of an infected human (*Ascaris lumbricoides*). Spread through the ingestion of eggs defected by an infected host, *A. lumbricoides* is suspected to infect over 700 million people worldwide[22,23]. The related species *A. suum* is considered one of the most prevalent helminths in domestic pigs [24]. These two closely related species are morphologically indistinguishable often capable of cross-transmission and sometimes hybridization with genetic introgression, and their taxonomic relationship (distinct or single shared species) is still debated [25–27]. Comparative genomic analyses demonstrate that the overall genomic divergence between *A. suum* and *A. lumbricoides* is minimal (<1%), driven primarily by single nucleotide polymorphisms (SNPs) [18,28,29].

First described in 1992 [30], the *Ascaris* mitochondrial genome has been well studied, enhancing our understanding of *Ascaris* population structure through the use of targeted individual markers. Among the most frequently used markers is the gene encoding *cox1* – a mitochondrial protein integral to oxidative phosphorylation. This has led to the identification of over 70 distinct haplotypes that cluster into three distinct clades (A-C) [31]. Clade A incorporates isolates from pigs in non-endemic regions and humans in endemic regions, while clade B includes mixed isolates from both pigs and humans which are geographically widespread irrespective of endemicity. In contrast, clade C predominantly comprises pig-derived isolates from Europe, and small number of human-derived isolates from non-endemic regions indicating potential zoonotic transmission [31]. This pig-specific clade forms a clear distinct cluster [28,31–33], which has been suggested to have diverged as recently as 2500 years ago [33–35]. Comparatively, clade A and B are suspected to have split more recently, as early as 300 years ago, and have been readily dispersed through international movement of people and pigs [34].

While *Ascaris* remains one of the most prevalent helminth infections globally, key aspects of its population biology and transmission dynamics are still poorly understood [27]. The close genetic similarity between *A. lumbricoides* and *A. suum*, coupled with evidence of hybridisation and introgression in regions where humans and pigs coexist, continues to obscure species boundaries and complicate the assessment of zoonotic potential [28,31,37,38]. Compared with many other nematodes, *Ascaris* nuclear genomes are notably large and range from 270–300 Mbp and contain approximately 18,000 protein-coding genes [39], posing substantial computational challenges for genome assembly and analysis. Population-level molecular studies have been further limited by the cost and complexity of sequencing and by reliance on single-locus markers with low discriminatory power, which offer insufficient resolution to infer population structure, gene flow, or the sources of reinfection following treatment. As a result, in-depth genomic data from endemic regions remain scarce, and the relative contribution of human and animal reservoirs to ongoing transmission is not well defined. Additionally, to achieve the WHO 2030 target, more detailed understanding of parasite population structure and transmission in endemic settings is essential to guide future diagnostic and control strategies.

Given that single-marker or reference-based approaches have often failed to fully resolve nematode population dynamics, *de novo* mitochondrial genome assemblies represent a valuable practical alternative. Beyond improving mitochondrial genome recovery, such workflows can be leveraged for high-throughput species identification and the detection of potential zoonotic reservoirs. Their implementation in endemic regions could further facilitate real-time epidemiological surveillance, inform integrated control programs, and enhance understanding of cross-species infection potential within the genus.

Here, we present a comprehensive workflow for reconstructing complete mitochondrial genomes from *Ascaris*, providing new insights into the population dynamics, evolutionary relationships, and host–parasite interactions of these globally important parasites.

## 2. METHODS

### 2.1 Sample information and initial processing

Twenty-four *Ascaris* worms obtained from humans and pigs from different countries worldwide were included in this study. Seven adult *Ascaris* samples from Germany were collected from slaughtered pigs at an abattoir located in the federal state of North Rhine-Westphalia (samples DE_P6-P24). Additionally, one adult worm was sent as diagnostic material to the Institute for Parasitology, University of Veterinary Medicine Hannover, from a human patient from the federal state of Schleswig-Holstein suffering from recurrent *Ascaris* spp. infection with suspected zoonotic transmission due to backyard pig farming (sample DE_H1a). Information on the origins of other new samples and downloaded mitogenomes is summarised in Supplementary Table S1. Ethical approvals for the original sample collections were obtained by the respective research groups and granted by the relevant institutional review boards and national authorities, as reported in the corresponding publications (see Table S1 for full details). All original studies were conducted in accordance with the appropriate institutional, national, and international guidelines and regulations.

In addition, Illumina sequencing reads from 125 *Ascaris* worms were downloaded from the NCBI Sequence Read Archive (https://www.ncbi.nlm.nih.gov/sra), while an additional 18 *Ascaris* mitogenomes and 1 *Toxascaris* mitogenome were obtained from NCBI Nucleotide (https://www.ncbi.nlm.nih.gov/nucleotide). This study did not involve the collection of new biological samples.

### 2.2 DNA extraction

DNA was extracted from ∼1-inch sections of worm tissue. To minimise contamination from fertilised eggs and male sperm (which could lead to spurious genotypes[40]), the uteruses and intestines were removed. Muscle tissue was separated from the cuticle and processed in-house using the Qiagen DNeasy Blood & Tissue Kit (Qiagen, Hilden, Germany; Cat. No. 69506) or the MasterPure Complete DNA and RNA Purification Kit (Lucigen, Middleton, WI, USA; Cat. No. MC85200) following the manufacturers’ protocols. For the Qiagen kit, DNA was eluted in 50 µl of AE buffer, which was re-applied to the spin column to maximise DNA concentration.

DNA concentration and purity were assessed using the Invitrogen Qubit^TM^ 1X dsDNA High Sensitivity quantification kit and NanoDrop spectrophotometer, respectively. Only samples with a concentration ≥ 5 ng/µl and A260/A280 ratio ≥ 1.6 were utilised for sequencing. Whole-genome sequencing was performed by Novogene Europe (Cambridge, UK) using the Illumina NovaSeq X Plus platform with 150 bp paired-end reads, at 30X coverage.

### 2.3 Processing genomic data

For all samples, raw sequence reads were processed with FastP (v0.23.4) [41] to remove any remaining adapter sequences and low-quality reads. Seqtk (version 1.5-r133)[42] was employed to subsample each paired-end read set to 20 million (20M) reads, totalling 40M reads per sample. Those samples which had less than 20M reads each way were not subsampled, but still utilised in the analysis. Quality control of the subsampled reads was assessed using FastQC (version v0.12.1) [43].

We compared whether host depletion or mtDNA enrichment improved the assemblies using Bowtie2 (Version 2.5.4) [44] (host depletion only) and Deacon (Version v0.7.0) [45,46]. For Bowtie2, a host reference database was constructed using Bowtie2-Build, incorporating the pig (GCF_000003025.6) and human (GCF_000001405.40) reference genomes, including their mitochondrial sequences. Subsampled 2× 20 million paired-end reads were aligned to deplete any reads mapping to the host genomes [47] using the SQM_mapper script version 1.65post1 [48]. A similar process was carried out for Deacon whereby a minimizer index file was created using the aforementioned human and pig reference genomes for depletion of host genome reads [45,46]. Deacon was also used for filtering *Ascaris* mtDNA reads using a database of mitogenomes from a variety of ascarids (see Supplementary Table S4 for full accession list). Only reads that were retained upon filtering were used for genome assembly.

*De novo* assembly of mitochondrial genomes was performed using MitoZ (Version v3.6) [49], initially employing *k*- of 39, 59, 79, 99, and 119 for all samples, except for those from Kenya [28], which had an average sequencing read length of 100 bp. Therefore, the *k-* of 119 was removed and the *k-* of 29, was added. Initially, 1 Gbp coverage was used for raw reads post-decontamination using Bowtie2 or Deacon, whereas 0.1 Gbp was utilised for the mtDNA alone. For those that failed assembly or produced partial or very large assemblies, further amendments to the *k* values (addition of k=21 or k=89) and lowered coverage requirements were used to achieve complete genome assembly. Sequence length and N50 for each assembly was assessed using QUAST (version 5.02) [50]. The workflow is depicted in Figure 1.

**Figure 1:**
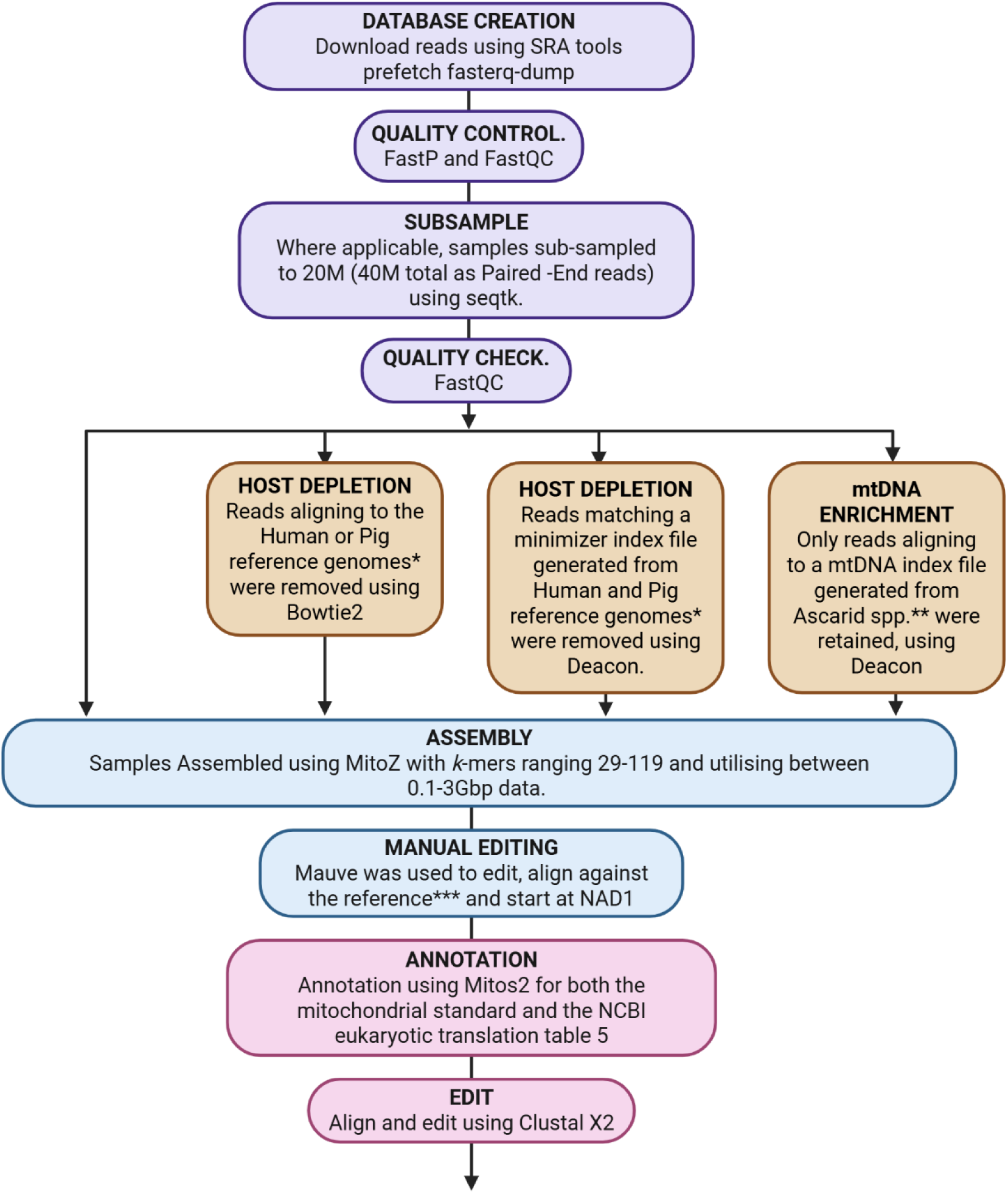
Pipeline for De Novo Assemble of mitogenomes from raw sequencing files. *GCF_000001405.40 and GCF_000003025.6, respectively. **See Supplementary Table S4 for full list. *** GCA_013433145.1

Each resulting FASTA file was then visually inspected using Mauve Aligner (version 2015-02-26) [51] to multi-align to the reference genome. When required, manual editing was performed to transform sequences into reverse complement and to linearise/align all sequences to start at the repetitive 5’ region ahead of the *nad1* gene. Annotation was performed using the standalone version of MitoS2 (version 2.1.9) [52] using the mitochondrial standard and NCBI translation table 5 (invertebrate mitochondrial code). Clustal X2 [53] was used to align and edit the resulting annotations.

Thirty samples—including all samples which were sequenced in this study and a subset of samples where whole genome sequence data was previously published (SRR31675200, SRR31675201, SRR31675202, SRR8419293, SRR8419294 and SRR8419295) — were aligned back to their respective assemblies to confirm that each assembly accurately represented the original sequencing reads. To achieve this, each assembly was indexed as a reference genome using Bowtie2-build, and the corresponding 20M subsampled reads were aligned back to their assemblies using Bowtie2 with the very-sensitive setting. Samtools (Version 1.22.1) [54] was used to ascertain coverage and average depth.

All assemblies were annotated using Mitos2 version 2.1.9 using codon translation table 5, the standard start/stop predictions and the NCBIcode start/stop predictions, with the NCBIcode prediction used for the *atp6*, *cox1*, *cox2*, *cox3*, *nad2*, *nad3* and *nad6* genes, and the standard start/stop prediction used for the *cob*, *nad1*, *nad4*, *nad4L* and *nad5* genes. For *COX1*, the annotation software missed the first 8 amino acids which were excluded from further analysis. All samples were oriented to begin with the repetitive region upstream of *NAD1* and were successfully annotated for all 12 protein-coding genes (Supplementary Figure S1).

We downloaded additional 19 complete mitochondrial genomes from GenBank encompassing both human and porcine hosts (n= 13), but also worms from non-human primates (n=2), sheep (*Ascaris ovis,* n=3) and one *Toxascaris leonina* sample (canine, China) to be used as an outgroup; the accession numbers are provided in Supplementary Table S1. This resulted in a final dataset of 168 samples.

### 2.4 Phylogenomic inference of *Ascaris* spp.

For both nucleotide and amino acid analyses, each annotated protein-coding gene was manually extracted into individual files and then concatenated in the order present in the assembly (*nad1–atp6–nad2–cob–cox3–nad4–cox1–cox2–nad3–nad5–nad6–nad4L*), excluding tRNA and rRNA genes, and repetitive OH region.

Neighbour-joining trees with bootstrap support (1,000 replicates) were constructed in MEGA12 [55] for the 12 individual gene alignments (nucleotide only) as well as for the 2 concatenated sequences. A neighbour-joining approach was chosen because it offers a computationally efficient means of visualising relationships among closely related mitochondrial genomes and has been widely applied in previous ascarid phylogenetic studies [32,56]. The resulting phylogenetic trees were visualised and analysed in R [57]. A heatmap illustrating clade assignment per gene for each sample, and plots demonstrating relationships among continent, host, and clade, and sample order rearrangement were created, also using R.

To assess if the resulting the assemblies retain the expected population-level variation, genetic diversity indices were calculated for each coding gene and concatenated dataset using DnaSP (version 5)[58], using the DNA polymorphism function.

## 3. RESULTS

### 3.1 Validation of the *de novo* mtDNA assembly pipeline of *Ascaris* spp

We generated Illumina sequence data from 24 *Ascaris* samples from pigs and humans, from varied geographical sources and combined them with downloaded Illumina fastq files from human samples from Kenya [28] and Ethiopia [59]. The final dataset included 134 samples from humans and 15 from pigs. Geographically, the samples represented three continents: Africa (Ethiopia [n=61], Uganda [n=7], Kenya [n=64]), Asia (Philippines [n=4], Thailand [n=3]), and Europe (Germany [n=8], Hungary [n=1], United Kingdom [n=1]).

Each set of paired end sequencing reads was subsampled to 2× 20 million reads and assembled with MitoZ either without further processing, depletion of pig and human reads with Bowtie2, depletion of pig and human reads with Deacon and filtering of mtDNA-specific reads with Deacon. Across four approaches, the vast majority of samples were successfully assembled (>95%), producing a single contig approximately 14 kb which contained all expected 12 protein-coding genes (Table 1, Supplementary Table S2). Partially complete assemblies were defined as those in which the full mtDNA was recovered but additional contigs were also produced, whereas incomplete assemblies generated contigs without resolving the full mtDNA, as defined by the presence of all 12 protein-coding genes.

**Table 1:** Assembly outcomes across unfiltered, host depletion, and mtDNA enrichment approaches. Assemblies were classified as *complete* when a single contig of the expected size and containing all 12 coding genes was produced; *partially complete* when full-length mtDNA was recovered but included additional contigs; *incomplete* when contigs were generated but no full mtDNA sequence was obtained; and *failed* when no assembly could be resolved.

|  | Subsampled<br>unfiltered data | Host depletion<br>using Bowtie2 | Host depletion<br>using Deacon | mtDNA-enrichment<br>with Deacon |
| --- | --- | --- | --- | --- |
| Mean contig length<br>(bp)* | 14,367 | 14,139 | 14,011 | 14,370 |
| No. of circular<br>assemblies (%) | 43/146<br>(29.5%) | 22/149<br>(14.8%) | 48/149<br>(32.2%) | 67/147<br>(45.6%) |
| No. complete<br>assemblies in one<br>contig. (%) | 142<br>(95.3%) | 145<br>(97.3%) | 146<br>(98.0%) | 142<br>(95.3%) |
| No. partially<br>complete assemblies<br>(%) | 1<br>(0.7%) | 1<br>(0.7%) | 1<br>(0.7%) | 1<br>(0.7%) |
| No. incomplete<br>assemblies (%) | 3<br>(2.0%) | 3<br>(2.0%) | 2<br>(1.3%) | 4<br>(2.7%) |
| No. failed assemblies<br>(%) | 3<br>(2.0%) | 0 | 0 | 2<br>(1.3%) |

\*Values for mean and median contig length are reported only for complete assemblies, and rounded to the nearest whole number.

Among completed assemblies, median contig lengths were highly consistent (13.98–14.37 kb), in line with the expected size of the *Ascaris* mitochondrial genome. Deacon depletion produced the shortest average contig (14,011bp), though values were comparable across all methods. Both host depletion strategies increased the proportion of successful assemblies compared with unfiltered or enriched datasets (∼97.5% vs 95.0%). Although mtDNA enrichment generated the highest number of circularised assemblies, it reduced the proportion of successful outcomes and yielded higher rates of incomplete and failed (2.0%) assemblies. Additionally, while the outcomes of both depletion approaches – including the average length of the complete assemblies – were in agreement, the number of circularised assemblies was far greater using Deacon compared with Bowtie2 (32.2% vs 14.8%, respectively). Notably, one sample (SRR31692075) was fully resolved in a single contig only with unfiltered data; other approaches either generated multiple contigs (depletion methods) or produced an abnormally large assembly (mtDNA enrichment).

Assemblies obtained following Bowtie2-based host depletion were designated as the reference approach. For datasets that could not be resolved using this method, the most complete alternative assembly (defined as the presence of all 12 coding genes) was selected for downstream analyses. Only one sample (SRR31692059) did not have a single contig assembly and required manual editing to resolve into a single contig, with a gap of 17 nucleotides based on alignment with other *Ascaris* mtDNA genomes, located in the intergenic region between the *cox1* and *cox2* genes.

To assess assembly quality and representativeness, the original sequencing reads were mapped back to their respective *de novo* mtDNA assemblies (see Supplementary Table S3). Of the 30 samples that underwent mapping, a very high coverage was achieved, with most samples exceeding 10,000 ×. The mean base depth ranged from 937× to 158,022×, while mean base quality scores ranged from 34.8 to 39.5, and average mapping quality was consistently high (≥41.4) across all samples. Importantly, all assemblies achieved >99.8% genome coverage.

### 3.2 Identification of a novel mtDNA clade for *Ascaris*

To investigate the genetic diversity and phylogenetic structure of *Ascaris* isolates, we constructed a neighbour joining tree using the concatenated mitochondrial protein-coding genes, the 12 individual genes and the translated concatenated amino acid sequence (Figures 2 and 3; Supplementary Figures S2-13). The resulting phylogeny revealed the presence of four well-supported mitochondrial clades, designated A, B, C based on previous assignment[34], and a newly identified clade D, as well as some potential sub-clades within both clades A and B. Consistent with previous studies [28,34], clade A comprised the majority of human-derived *Ascaris* samples, forming a large, well-supported cluster with high bootstrap values (100%). Clade B represented a more mixed host population, including both human and non-human hosts, with both clades A and B containing geographically mixed samples.

**Figure 2.**
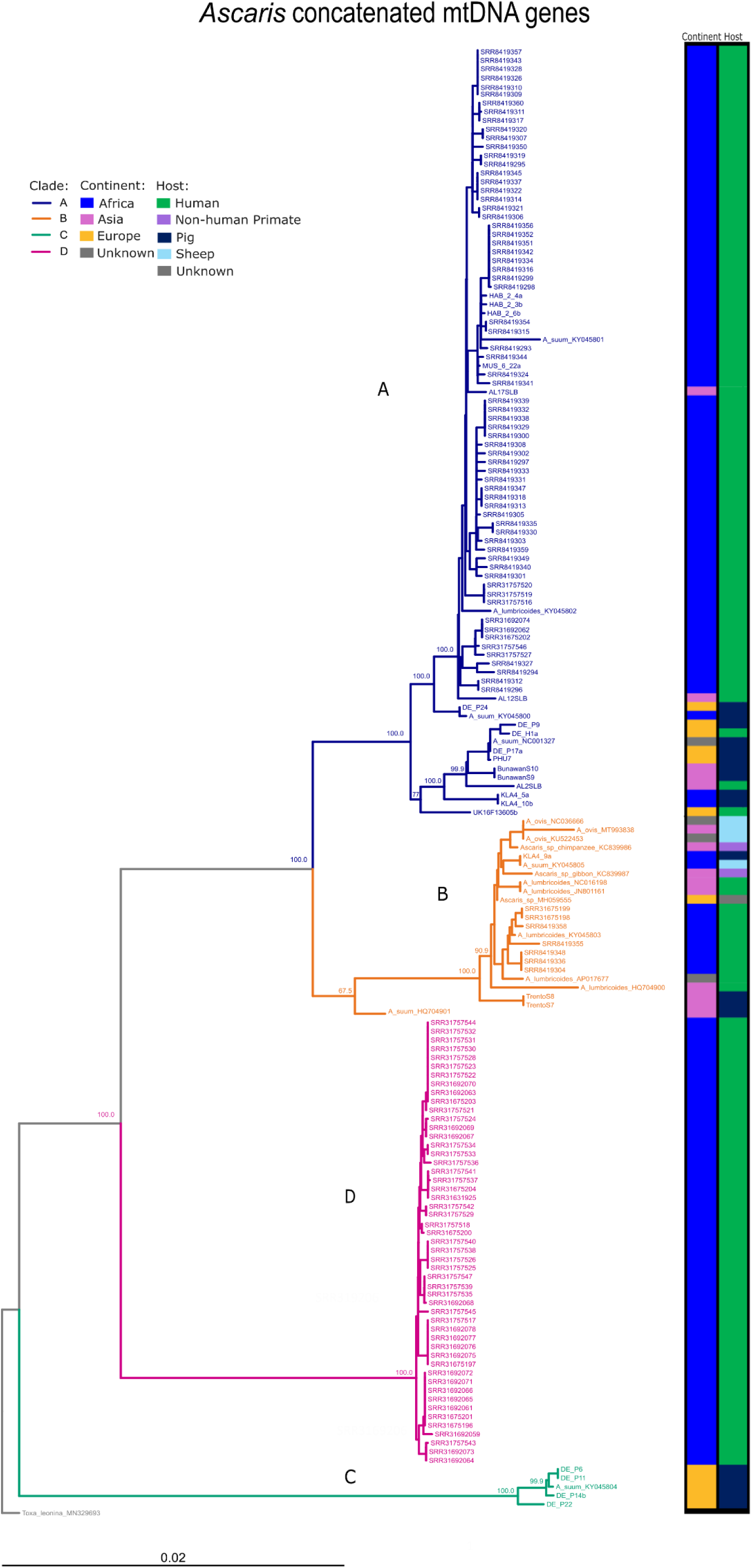
Neighbour-joining tree of concatenated mitochondrial DNA sequences from Ascaris samples. The tree was constructed using concatenated nucleotide sequences from 12 mitochondrial protein-coding genes in the order: *nad1–atp6–nad2–cob–cox3–nad4–cox1–cox2–nad3–nad5–nad6–nad4L*. Bootstrap values (>80%) from 1,000 replicates are shown at major nodes. Tip labels correspond to individual sample identifiers. The tree is drawn to scale, with branch lengths in the same units as those of the evolutionary distances to infer the phylogenomic tree. The evolutionary distances were computed using the Maximum Composite Likelihood Method and are in the units of the number of base substitutions per site. The analysis encompassed 168 sequences, representing 10,247 positions. Analysis was conducted using MEGA12. Clades A–D were defined based on phylogenetic clustering. Colour indicates clade assignment as follows: blue (A), orange (B), green (C), and pink (D). The tree is rooted using the outgroup *Toxascaris leonina* MN329693 (scale bar: 0.02, Optimal tree with the sum of branch length =0.308). To the right of the tree, two annotation columns indicate the continent (inner bar) and host species (outer bar) associated with each sample. Continent (Africa (red), Asia (orange), Europe (green), Unknown (grey)) and host (Human (blue), Pig (cyan), Sheep (purple), Non-human Primate (navy), Unknown (dark grey)) are colour-coded as shown in the legend.

**Figure 3:**
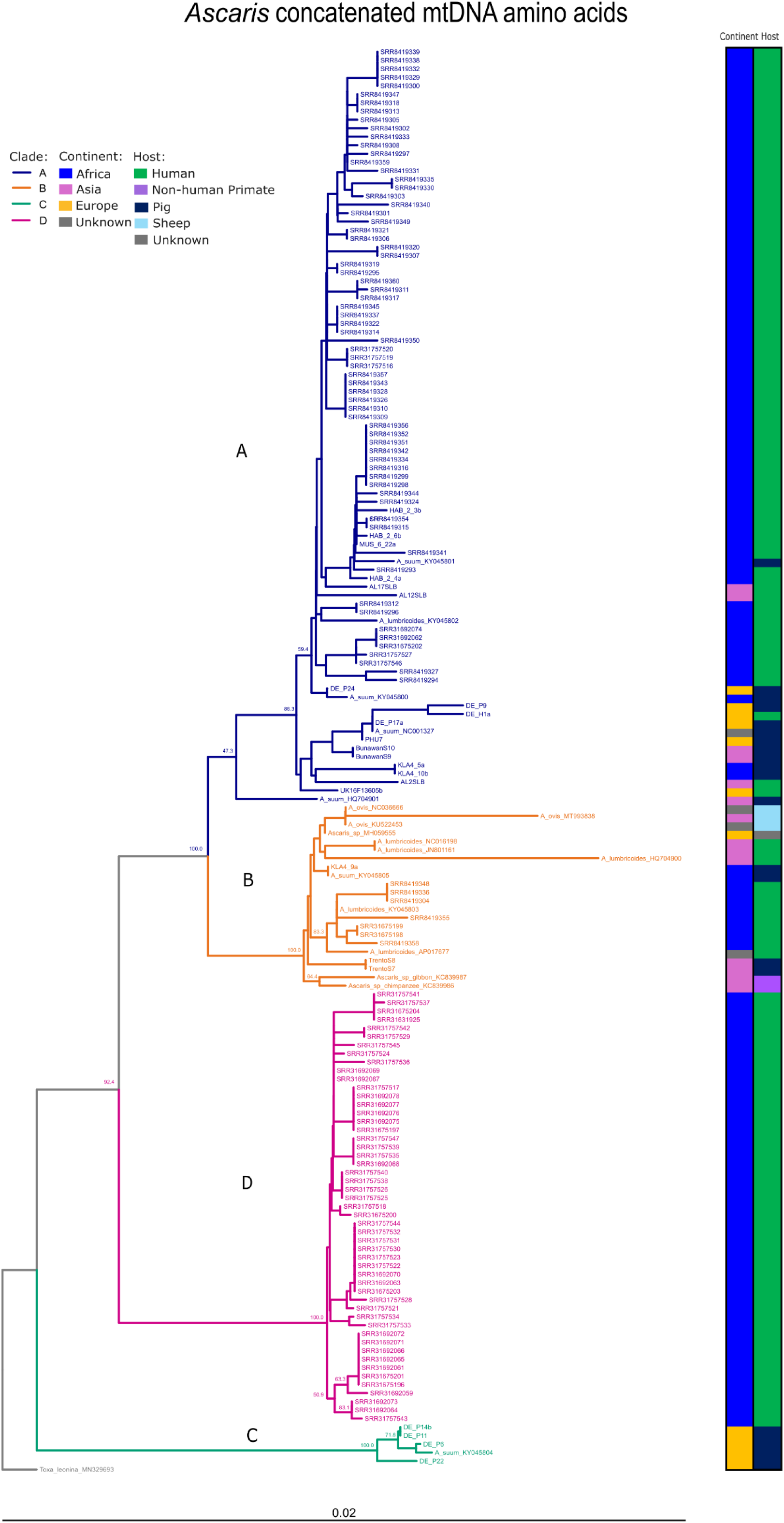
Neighbour-joining tree of concatenated mitochondrial amino acid sequences from Ascaris samples. The tree was constructed using concatenated amino acid sequences from 12 mitochondrial protein-coding genes in the order: *nad1–atp6–nad2–cob–cox3–nad4–cox1–cox2–nad3–nad5–nad6–nad4L*. Bootstrap values from 1,000 replicates are shown at major nodes. Evolutionary distances are inferred using the Poisson correction method and are in the unit of amino acid substitution per site. The pairwise deletion option was applied to all the ambiguous positions for each sequence resulting in a final dataset comprising of 3,405 positions. Analysis was conducted using MEGA12. Samples were grouped into major clades (A–D) based on phylogenetic topology and bootstrap support. Colour indicates clade assignment as follows: blue (A), orange (B), green (C), and pink (D). The tree is rooted using the outgroup *Toxascaris leonina* MN329693 (scale bar: 0.02 nucleic acid substitutions per site). To the right of the tree, two annotation columns indicate the continent (inner bar) and host species (outer bar) associated with each sample. Continent (Africa (red), Asia (orange), Europe (green), Unknown (grey)) and host (Human (blue), Pig (cyan), Sheep (purple), Non-human Primate (navy), Unknown (dark grey)) are colour-coded as shown in the legend.

We identified further samples which were assigned to Clade C, previously identified as comprising *Ascaris* samples from pigs in Europe, from our pool of *Ascaris* samples from Germany. However, the human infection case (DE_H1) which was associated with close contact with pigs via backyard farming practices, along with a subset of pig-derived worms (n = 3) from a German abattoir, clustered within Clade A, rather than Clade C.

The novel clade D presented as a genetically distinct and well-supported group, suggesting the presence of a divergent lineage or cryptic population. To our knowledge this is the first time this clade has been described, where all samples within this clade were derived from human hosts in Ethiopia, collected as part of a community-wide deworming initiative [59]. A small number of the samples from Ethiopia (n = 10, 16.4%) were assigned to Clades A (n = 8) and B (n = 2), indicating the presence of multiple mitochondrial lineages circulating within a single geographical setting.

Most samples analysed were derived from African populations, predominantly from human hosts, leading to an overrepresentation of African *Ascaris* isolates in the dataset. Despite broad geographical distribution, mitochondrial clades were not uniquely separated by continent or host; however some structuring was evident. In particular, clade C, which contains primarily samples obtained from pigs from Europe (Figure 4).

**Figure 4.**
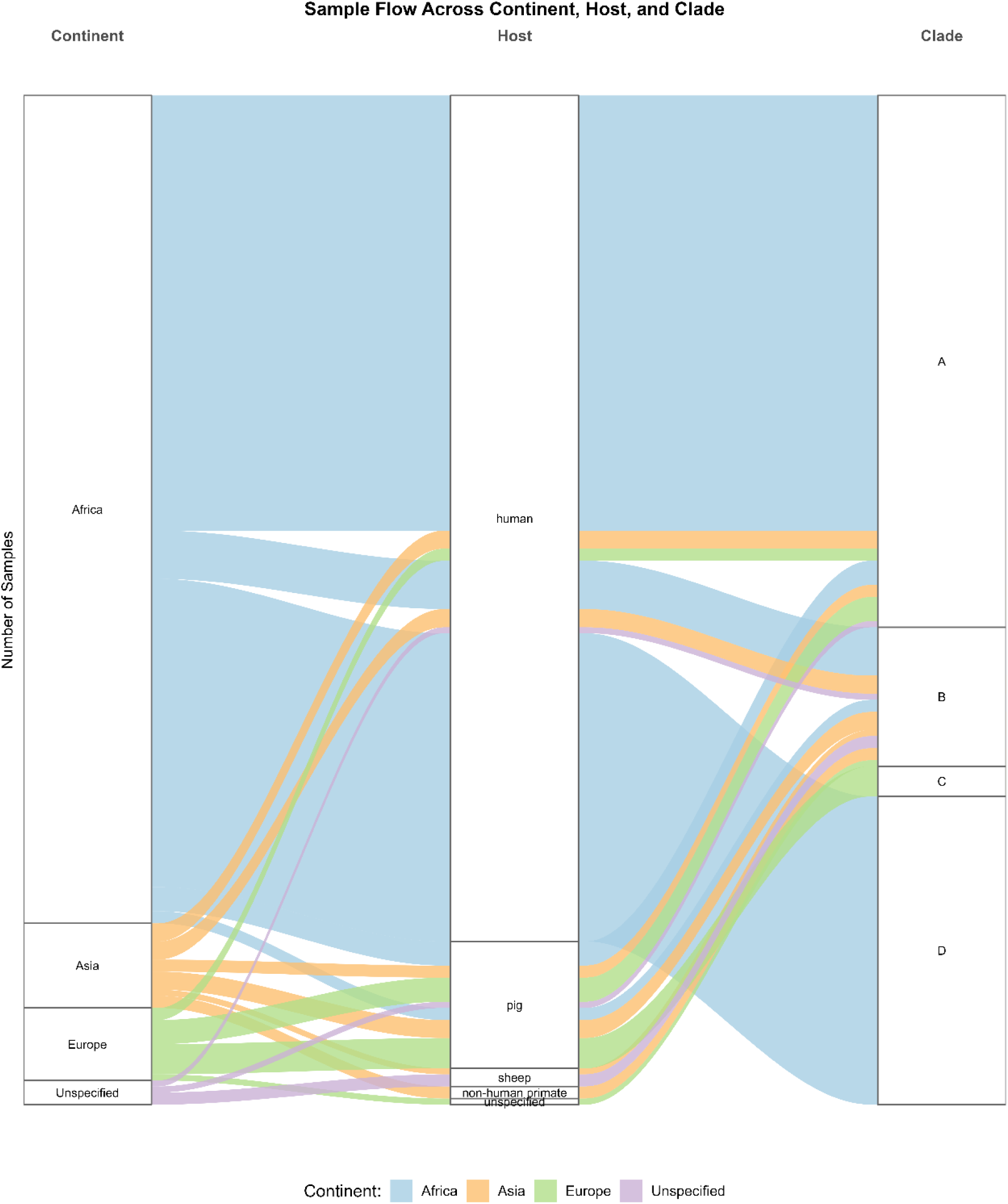
Alluvial plot illustrating the flow of mitochondrial Ascaris samples across continent of origin, host species, and mitochondrial clade (collapsed to A–D). Stratum widths and flow thicknesses are proportional to the number of samples within each category. The majority of samples originated from humans in Africa and were assigned to clades A or D. Colour represents continent of origin. The plot highlights distinct geographical and host-specific structuring across clades.

We observed strong concordance between phylogenetic placements based on concatenated nucleotide sequences and those based on translated amino acid alignments. Most samples retained consistent positions across both trees, with only minor within-clade rearrangements. The major clades (A–D) remained well-supported and topologically stable between datasets, indicating that both nucleotide and amino acid data capture similar evolutionary signals. To visualise these differences, samples were plotted as a tanglegram in the order as they appeared in each tree, with connecting lines indicating positional shifts (Figure 5). Crossing lines indicate discordance in sample placement, which in most cases was limited to within-clade shifts. The only sample where this did not apply, was sample HQ704901 (pig, China) [60] which shifted from clade B to clade A between the nuclear and amino acid trees, respectively.

**Figure 5:**
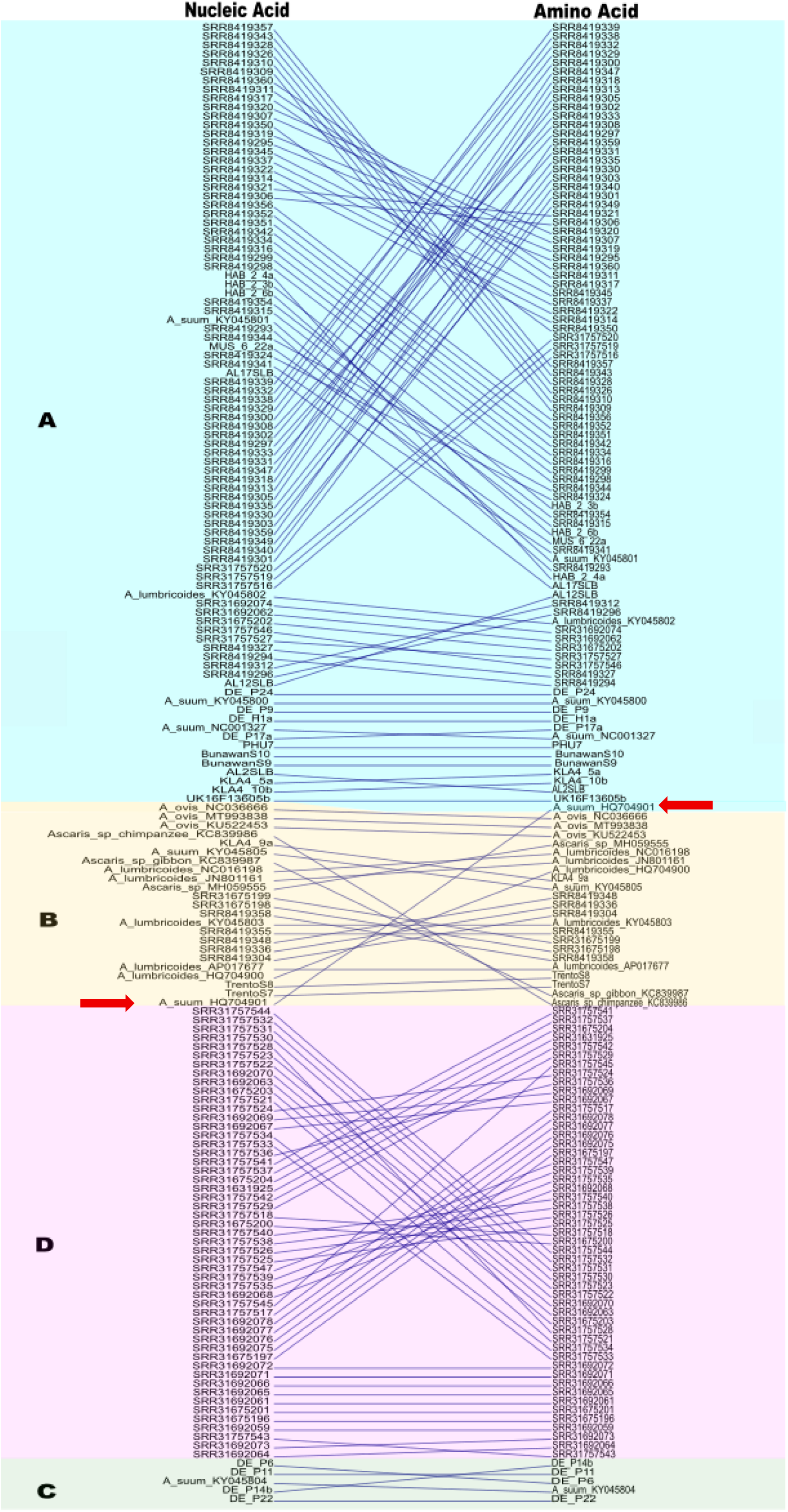
Tanglegram showing sample order rearrangement between nucleic acid and amino acid phylogenetic trees for 168 Ascaris mtDNA assemblies. Each line represents a single sample and connects its position in the nucleotide-based tree (left) to its corresponding position in the amino acid-based tree (right). Samples are coloured by their tree clade membership (A–D). Crossing lines primarily indicate within-clade reordering, or for the one mtDNA-discordance between the two tree topologies sample HQ704901 [60](as indicated by the red arrows)

### 3.3. Gene-wise and Concatenated Mitochondrial Analyses of Phylogenetic Signal and Diversity

To further evaluate whether any single mitochondrial gene contributed disproportionately to the phylogenetic positions, we generated neighbourhood-joining trees for each individual coding gene (Supplementary Figures S2-13). For each gene, clade membership was determined per sample based on tree visualisation and bootstrap support. The resulting clade associations were compiled into a summary heatmap (Figure 5), which was largely concordant across individual genes and consistent with the concatenated datasets. An exception was observed in the gene *nad4L*, a relatively short (∼234bp in length [60]), highly conserved gene (Table 2), which clustered clades A and B together. Additionally, one notable outlier, the aforementioned sample HQ704901 [60], showed significant incongruence. As previously reported [34] HQ704901 did not cluster clearly with either clade A or B. The per-coding-gene analysis supports this observation: 7 of the mitochondrial genes assigned HQ704901 to clade A, while the remaining 5 genes clustered with clade B (Figure 6). This phylogenetic ambiguity is also reflected in the concatenated trees, where HQ704901 forms a distant branch nested within the broader clade B lineage for nucleic acids and A for amino acids.

**Figure 6.**
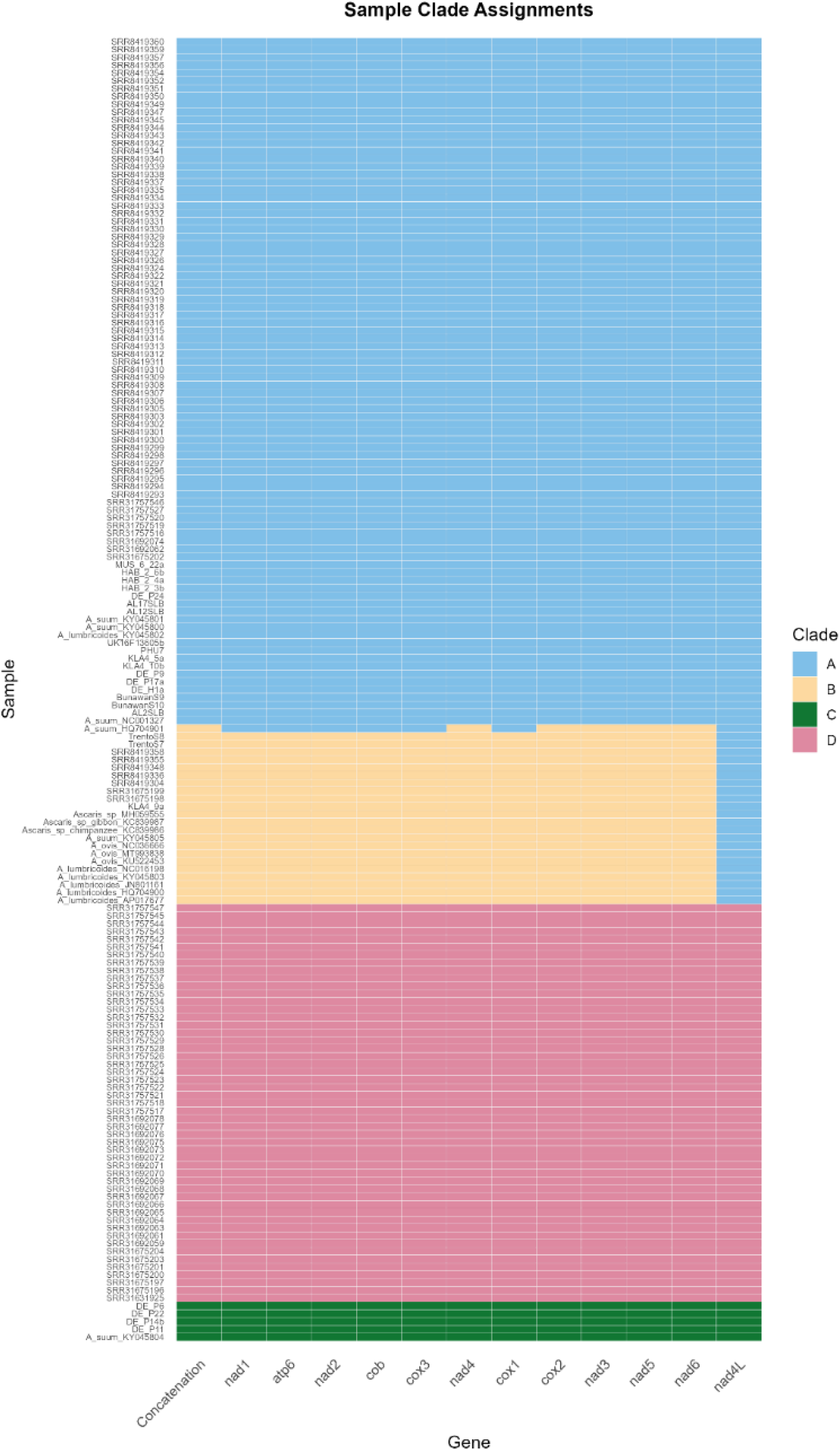
Heatmap of clade assignment for each mitochondrial gene across all Ascaris samples. Each row represents a sample, and each column corresponds to a mitochondrial gene, including both individual protein-coding genes and the full concatenated sequence. Colours denote clade assignment per gene, as inferred from phylogenetic analysis. Blue, orange, pink, and green represent clades A, B, D, and C, respectively. This visualisation highlights the consistency (or discordance) of clade assignment across genes and samples. Most samples show concordant assignment across all genes, while a minority exhibit inter-gene variability, reflecting gene-specific phylogenetic signal or possible recombination.

**Table 2:** Mitochondrial genetic diversity across reconstructed assemblies.

| Gene / Region | Length (bp)* | Segregating sites (S) | S/kb ((S/ Length (bp)*1000) | Haplotypes (h) | Haplotype Diversity (±SD) | $\pi$ (±SD) | $\theta_W$ | K |
| --- | --- | --- | --- | --- | --- | --- | --- | --- |
| Concatenated dataset | 10240 | 1261 | 123.2 | 94 | 0.987 (0.003) | 0.0224 (0.0010) | 0.0216 | 228.9 |
| *nad1* | 872 | 130 | 149.1 | 44 | 0.938 (0.009) | 0.0239 (0.0010) | 0.0262 | 20.795 |
| *atp6* | 600 | 75 | 125.0 | 44 | 0.897 (0.015) | 0.0177 (0.0009) | 0.022 | 10.594 |
| *nad2* | 843 | 108 | 128.1 | 43 | 0.862 (0.020) | 0.0218 (0.0011) | 0.0225 | 18.384 |
| *cob* | 1096 | 141 | 128.7 | 41 | 0.912 (0.013) | 0.0202 (0.0011) | 0.0226 | 22.151 |
| *cox3* | 777 | 97 | 124.8 | 44 | 0.883 (0.016) | 0.0219 (0.0011) | 0.0219 | 17.009 |
| *nad4* | 1230 | 188 | 152.8 | 55 | 0.954 (0.007) | 0.0258 (0.0013) | 0.0269 | 31.785 |
| *cox1* | 1534 | 151 | 98.4 | 69 | 0.966 (0.007) | 0.0208 (0.0008) | 0.0173 | 31.224 |
| *cox2* | 699 | 67 | 95.9 | 33 | 0.839 (0.019) | 0.0171 (0.0009) | 0.0168 | 11.979 |
| *nad3* | 336 | 34 | 101.2 | 21 | 0.814 (0.020) | 0.0232 (0.0010) | 0.0178 | 7.78 |
| *nad5* | 1584 | 188 | 118.7 | 61 | 0.963 (0.007) | 0.0257 (0.0011) | 0.0209 | 40.635 |
| *nad6* | 435 | 54 | 124.1 | 29 | 0.814 (0.021) | 0.0275 (0.0012) | 0.0218 | 11.967 |
| *nad4L* | 234 | 28 | 119.7 | 19 | 0.801 (0.018) | 0.0197 (0.0014) | 0.021 | 4.608 |

\* Excluding gaps and missing data
Abbreviations: S = number of segregating sites; h = number of haplotypes; $\pi$ = nucleotide diversity per site; $\theta_W$ = Watterson's estimator of $\theta$ (estimation the population mutation rate ( $\theta$ ), based on the number of segregating sites (S) in a sample); K = average number of pairwise nucleotide differences; SD = standard deviation.

To evaluate the performance of the assemblies for downstream population genetic analyses, we examined genetic diversity across the mitochondrial genome. All mtDNA assemblies were used to estimate genetic diversity indices across 12 protein-coding genes and the concatenated dataset (Table 2). The number of segregating sites per kb ranged from 95.9 (*cox2*) to 152.8 (*nad4*), demonstrating reliable recovery of variation across genes of varying size and both haplotype and nucleotide diversity (π) values were consistently high. Additionally, estimates of nucleotide diversity (π) and Watterson’s estimator of the population mutation rate (θ), calculated from the number of segregating sites (S) (θW), were broadly concordant, indicating unbiased detection of variation. The *nad4*, *nad5* and *nad6* genes exhibited the highest nucleotide diversity, with *nad4* and *nad5* displaying the greatest average number of pairwise differences (K). The concatenated dataset captured over 1,200 segregating sites and 94 haplotypes, confirming that the reconstructed assemblies retain sufficient resolution to recover population-level diversity across the mitochondrial genome.

## 4. DISCUSSION

The harnessing of genomic data is becoming increasingly central to public health surveillance, via the monitoring of parasite transmission, the discovery of hotspots of focal infection and identification of zoonotic or geographically driven transmission routes. In this study, we aimed to enhance the assembly of *Ascaris* mitochondrial genomes from low-coverage whole genome sequence data. Although some minor manual curation was still required, the majority of assemblies were developed using freely available, previously validated bioinformatic tools, supporting reproducibility and long-term sustainability.

The removal of host DNA bioinformatically, otherwise known as host sequence depletion, is becoming increasingly utilised in genomic analyses [61,62]. Several approaches for host decontamination have been successfully implemented, including retention of reads that align to a target reference genome [28,44,63], removal of reads aligning to the host genome [44,64,65] and utilisation of *k*-mers to filter out those specific to the host or utilising *k*-mer profiles for taxonomic classification[46,66].

Across the host sequence depletion and mtDNA filtering methods tested in this study, median complete contig lengths were highly consistent across methods (13.98–14.36 kb), in line with the expected size of the *Ascaris* mitochondrial genome. Assemblies generated using Deacon [46] depletion were consistently marginally shorter than those obtained through other approaches. This may reflect an improved removal of host or non-target reads and may be why circularisation was more frequent compared with bowtie2. Overall, host sequence depletion using either program resulted in superior assembly outcomes. It is important to note, however, that although the total number of assemblies was similar across both methods, the incomplete or partially complete samples did not always overlap. For instance, sample SRR31675203 was successfully resolved only through straight assembly and Bowtie2 depletion, whereas sample DE_P24 remained incomplete solely under Bowtie2 depletion. Furthermore, Deacon is a recently developed tool that offers significantly faster performance, particularly on lower-powered computational systems. Its continuous updates in response to user feedback enhance its functionality and usability, making it especially well-suited for large-scale analyses and resource-limited environments. These features position Deacon as a promising alternative to Bowtie2 for future applications.

Previous efforts to reconstruct *Ascaris* mitochondrial genomes from whole genome sequences have relied on reference-guided strategies requiring error correction and the use of GapFiller to span fragmented scaffolds [28]. This agrees with our data, where we have shown how mtDNA enrichment led to an increased number of contigs of variable length, and performed similarly to utilising the unfiltered reads alone. However, reference-guided approaches risk masking novel variation and may obscure population structure at a local and international level [67,68]. In *Ascaris*, this is compounded by the limited availability of high-quality reference genomes that span both human and porcine-derived *Ascaris* species and adequately represent the global distribution of the parasite [69]. Alternative approaches for mitochondrial genome reconstruction include long-range PCR and hybridisation capture to enrich *Ascaris* mtDNA [70]. Both of which can improve recovery but introduce distinct limitations. PCR-based methods are prone to amplification bias and may fail to capture the full extent of genetic diversity, while hybridisation capture, though proven very effective (six-fold enhancement), is costly and requires additional laboratory optimisation [70].

Our assembly workflow yielded mean alignment rates exceeding 98% to the annotated mitogenome and produced single-scaffold assemblies for over 98% of samples. Individual gene lengths were comparable to prior annotations [29,71], and estimates of haplotype and nucleotide diversity were within expected ranges, supporting the accuracy of the assemblies and their suitability for downstream phylogenomic and population genomic applications.

The phylogenies inferred from both individual gene trees and concatenated datasets revealed highly similar topologies, with clade C consistently distinct from the more closely related clades A and B, aligning with previous findings [31–33,72,73]. Where single genes often recover the same major clades, we have identified how they may miss signals of introgression or hybridisation and fail to resolve finer-scale structure/subclades. For instance, sample HQ704901 falls into the same clade when using standard mitochondrial markers including, *cox1* and *nad1.* However, *nad4*, despite having the highest mutation density per length, produces a different topology [34]. Thus, reliance on single-gene markers alone can obscure complex evolutionary events and fine-scale population structure.

A notable finding was the identification of a new clade D, which appears as an intermediate between clade C and the clades A and B, suggesting it may represent a transitional lineage, historical gene flow or incomplete lineage sorting. These samples were obtained from a single community in Ethiopia, collected as part of a longitudinal epidemiological study, where genomic analyses previously demonstrated fine-scale population structure with localised transmission predominantly at the household level rather than random community-wide mixing [59]. Despite an overall reduction in worm populations, likely resulting from mass drug administration efforts, a minority of human hosts continued to harbour disproportionately high worm burdens, which may explain the presence of a small number of worms in clades A and B, where differences in treatment coverage and compliance, multiple introduction events, and household-level micro-environmental factors may allow these ancestral lineages to persist [59].

In agreement with previous studies, we observed the same clade separation between worm populations infecting humans and those infecting pigs [19,31,74] (Figure 4). For example, we further identified European samples, in our case from German porcine hosts, in Cluster C. Geographical structuring was evident in our dataset; however, we do note that there is a significant bias towards samples derived from humans in Africa.

Previous studies have demonstrated that mitochondrial DNA differentiates between both geographic and species-level variation, as shown in Tanzanian [75] and Chinese[71] *Ascaris* populations. However, the whole genome sequence of Kenyan isolates indicated extensive interbreeding and a shared nuclear gene pool, despite divergent mitochondrial lineages[28] and nuclear microsatellite data from the same Tanzanian population suggested high gene flow and limited geographic structure [75]. Data from microsatellite analysis have also been interpreted to support either a single host shift followed by geographical separation [31] and initial geographical separation followed by multiple host shifts [38]. As microsatellites capture more recent evolutionary events compared with mtDNA, these contrasting signals may be clarified with broader sampling and increased availability of whole genome sequence data across hosts and regions. While incomplete lineage sorting of mitochondrial genomes may also contribute to the discordant phylogenetic patterns observed, their ability to capture deeper evolutionary events ensures they remain valuable for detecting broad-scale host- and geography-associated patterns.

In conclusion, we present a robust and scalable framework for the efficient recovery of high-quality mitochondrial genomes from low-coverage whole-genome sequencing datasets, thereby reducing reliance on targeted sequencing approaches. The integration of freely available bioinformatic tools provides for a flexible framework that can be readily adapted for large-scale analyses. By overcoming common challenges associated with fragmented or low-input samples, this approach enabled the discovery of a novel mtDNA clade for *Ascaris* and facilitated investigation into whether *A. suum* and *A. lumbricoides* represent distinct species or host-adapted populations within a single interbreeding complex. The setup provides a valuable resource for expanding genomic coverage across underrepresented populations and for supporting studies of parasite evolution, host specificity, and transmission dynamics. Together, these findings highlight the importance of integrating genomic surveillance and standardized preservation strategies into parasitic disease control frameworks, enabling more accurate monitoring of transmission dynamics and supporting global efforts toward the elimination of soil-transmitted helminths.

## 5. Data availability

All code is available at: https://github.com/LWoolfe/Ascaris_Mitogenome_Project.git. This includes all bioinformatic tools and R packages. The mtDNA assemblies have been deposited in the European Nucleotide Archive (ENA) as Bioproject PRJEB105081, with assembly accession numbers available via the Github page. The assemblies and the alignments of the 12 individual protein-coding genes and the concatenated genes and corresponding amino acid sequences have been made available via Figshare under the DOI: 10.6084/m9.figshare.30871202

## 6. Acknowledgements

We gratefully acknowledge the individuals and laboratories who generously donated worms used in this study.

## 7. Additional Information

### 7.2 Competing Interest Statement The authors declare no competing interests

#### 7.3 Authorship contribution

**LW** – Conceptualisation, Methodology, Investigation, Formal analysis, Data curation, Writing – Original Draft, Visualisation.

**K.K, U.C, K.K.L, A.J.A, P.A, A.J, J.R.S, C.S, M.K.R, S.P.L, T.L, V.G.P** – Resources, Investigation, Writing – Review & Editing.

**A.H.M.vV** – Methodology, Software, Formal analysis, Supervision, Writing – Review & Editing.

**M.B** - Supervision, Resources, Writing – Review & Editing.

