## Supplementary Information (Tables S1-S4, Figures S1-S13 for "Scalable Assembly of *Ascaris* Mitogenomes from Whole-Genome Data Reveals a Novel Clade"

Contains:

Supplementary Tables S1-S4

Supplementary Figures S1-S13

**Supplementary Table S1: Sample Information**

| Country | Host | Sample | Named species | Sequence data source | Reference for sample/data |
|---|---|---|---|---|---|
| **China** | Human | **HQ704900** | *A. lumbricoides* | Downloaded from NCBI Genome | [1] |
| **China** | Pig | **HQ704901** | *A.suum* | Downloaded from NCBI Genome | [1] |
| **China** | Sheep | **MT993838** | *A. ovis* | Downloaded from NCBI Genome | [2] |
| **China** | Non-Human Primate | **KC839986** | *Ascaris* | Downloaded from NCBI Genome | [3] |
| **China** | Non-Human Primate | **KC839987** | *Ascaris* | Downloaded from NCBI Genome | [3] |
| **Denmark** | Unspecified | **MH059555** | *Ascaris* | Downloaded from NCBI Genome | [4] |
| **Ethiopia** | Human | **SRR31631925 -SRR31675204; SRR31692059; SRR31692061 - SRR31692078; SRR31757516 - SRR31757547** | *A. lumbricoides* | Downloaded from SRA | [5] |
| **Germany** | Human | **DE_H1a** | *Ascaris* | This study | This study |
| **Germany** | Pig | **DE_P11** | *Ascaris* | This study | This study |
| **Germany** | Pig | **DE_P14b** | *Ascaris* | This study | This study |
| **Germany** | Pig | **DE_P17a** | *Ascaris* | This study | This study |
| **Germany** | Pig | **DE_P22** | *Ascaris* | This study | This study |
| **Germany** | Pig | **DE_P24** | *Ascaris* | This study | This study |
| **Germany** | Pig | **DE_P6** | *Ascaris* | This study | This study |
| **Germany** | Pig | **DE_P9** | *Ascaris* | This study | This study |
| **Hungary** | Pig | **PHU7** | | This study | [6] |
| **Kenya** | Human | **SRR8419293 - SRR8419324; SRR8419324; SRR8419326 - SRR8419336; SRR8419338 - SRR8419345; SRR8419347 - SRR8419360** | *A. lumbricoides* | Downloaded from SRA | [7] |
| **Korea** | Human | **JN801161** | *A. lumbricoides* | Downloaded from NCBI Genome | [8] |
| **Korea** | Human | **NC016198** | *A. lumbricoides* | Downloaded from NCBI Genome | [8] |
| **Philippines** | Pig | **Bunawan S9** | *Ascaris* | This study | [6] |
| **Philippines** | Pig | **Bunawan S10** | *Ascaris* | This study | [6] |
| **Philippines** | Pig | **TrentoS7** | *Ascaris* | This study | [6] |
| **Philippines** | Pig | **TrentoS8** | *Ascaris* | This study | [6] |
| **Tanzania** | Pig | **KY045800** | *A.suum* | Downloaded from NCBI Genome | [9] |
| **Tanzania** | Human | **KY045802** | *A. lumbricoides* | Downloaded from NCBI Genome | [9] |
| **Tanzania** | Human | **KY045803** | *A. lumbricoides* | Downloaded from NCBI Genome | [9] |
| **Thailand** | Human | **AL12SLB** | *Ascaris* | This study | [10] |
| **Thailand** | Human | **AL17SLB** | *Ascaris* | This study | [10] |
| **Thailand** | Human | **AL2SLB** | *Ascaris* | This study | [10] |
| **Uganda** | Human | **HAB_2_3b** | *Ascaris* | This study | [11] |
| **Uganda** | Human | **HAB_2_4a** | *Ascaris* | This study | [11] |

| | | | | | |
|---|---|---|---|---|---|
| **Uganda** | Human | **HAB_2_6b** | *Ascaris* | This study | [11] |
| **Uganda** | Pig | **KLA4_10b** | *Ascaris* | This study | [12] |
| **Uganda** | Pig | **KLA4_5a** | *Ascaris* | This study | [12] |
| **Uganda** | Pig | **KLA4_9a** | *Ascaris* | This study | [12] |
| **Uganda** | Pig | **KY045801** | *A.suum* | Downloaded from NCBI Genome | [9] |
| **Uganda** | Pig | **KY045805** | *A.suum* | Downloaded from NCBI Genome | [9] |
| **Uganda** | Human | **MUS_6_22a** | *Ascaris* | This study | [11] |
| **United Kingdom** | Pig | **KY045804** | *A.suum* | Downloaded from NCBI Genome | [9] |
| **United Kingdom** | Human | **UK16F13605b** | *Ascaris* | This study | [12] |
| **Unspecified** | Human | **AP017677** | *A. lumbricoides* | Downloaded from NCBI Genome | Unpublished |
| **Unspecified** | Sheep | **KU522453** | *A. ovis* | Downloaded from NCBI Genome | Unpublished |
| **Unspecified** | Pig | **NC001327** | *A. suum* | Downloaded from NCBI Genome | [13] |
| **Unspecified** | Sheep | **NC036666** | *A. ovis* | Downloaded from NCBI Genome | Unpublished |
| **China** | Dog | **MN329693** | *Toxascaris leonina* | Downloaded from NCBI Genome | [14] |

**Supplementary Table S2: mtDNA Assembly results after depleting host (human+pig) reads or enriching for mtDNA reads**

| | 20M unfiltered | | | | | bowtie2-filtered | | | | | deacon-filtered | | | | | mtDNA-enriched | | | | |
|---|---|---|---|---|---|---|---|---|---|---|---|---|---|---|---|---|---|---|---|---|
| **Sample** | contigs | mitocontig | Total | N50 | circular | contigs | mitocontig | Total | N50 | circular | contigs | mitocontig | Total | N50 | circular | # contigs | mitocontig | Total | N50 | circular |
| AL12SLB | 1 | 14433 | 14433 | 14433 | no | 1 | 13981 | 13981 | 13981 | no | 1 | 13911 | 13911 | 13911 | no | 2 | 8896 | 14042 | 8896 | no |
| AL17SLB | 1 | 14270 | 14270 | 14270 | no | 1 | 14388 | 14388 | 14388 | no | 1 | 13960 | 13960 | 13960 | no | 1 | 14306 | 14306 | 14306 | yes |
| AL2SLB | 1 | 14947 | 14947 | 14947 | no | 1 | 14262 | 14262 | 14262 | no | 1 | 14262 | 14262 | 14262 | no | 1 | 14430 | 14430 | 14430 | no |
| BunawanS10 | 1 | 14595 | 14595 | 14595 | no | 1 | 14300 | 14300 | 14300 | yes | 1 | 14003 | 14003 | 14003 | no | 1 | 14304 | 14304 | 14304 | yes |
| BunawanS9 | 2 | 14578 | 31149 | 16571 | no | 1 | 14085 | 14085 | 14085 | no | 1 | 14369 | 14369 | 14369 | no | 1 | 14308 | 14308 | 14308 | yes |
| DE_H1a | 1 | 14437 | 14437 | 14437 | no | 1 | 14429 | 14429 | 14429 | no | 1 | 13885 | 13885 | 13885 | no | 1 | 14519 | 14519 | 14519 | yes |
| DE_P11 | 1 | 14186 | 14186 | 14186 | yes | 1 | 14089 | 14089 | 14089 | no | 1 | 14075 | 14075 | 14075 | no | 1 | 14186 | 14186 | 14186 | yes |
| DE_P14b | 1 | 14027 | 14027 | 14027 | yes | 1 | 14027 | 14027 | 14027 | yes | 1 | 14024 | 14024 | 14024 | yes | 1 | 14160 | 14160 | 14160 | yes |
| DE_P17a | 1 | 15614 | 15614 | 15614 | no | 1 | 13844 | 13844 | 13844 | no | 1 | 13833 | 13833 | 13833 | no | 1 | 14402 | 14402 | 14402 | yes |
| DE_P22 | 1 | 14186 | 14186 | 14186 | yes | 1 | 14014 | 14014 | 14014 | no | 1 | 14137 | 14137 | 14137 | no | 1 | 14515 | 14515 | 14515 | no |
| DE_P24 | 1 | 14890 | 14890 | 14890 | no | 1 | 13319 | 13319 | 13319 | no | 1 | 13967 | 13967 | 13967 | no | 1 | 14330 | 14330 | 14330 | yes |
| DE_P6 | 1 | 14297 | 14297 | 14297 | yes | 1 | 14082 | 14082 | 14082 | no | 1 | 13905 | 13905 | 13905 | no | 1 | 14227 | 14227 | 14227 | no |
| DE_P9 | 1 | 14297 | 14297 | 14297 | yes | 1 | 14418 | 14418 | 14418 | yes | 1 | 14141 | 14141 | 14141 | yes | 1 | 14299 | 14299 | 14299 | yes |
| HAB_2_3b | 1 | 14728 | 14728 | 14728 | no | 1 | 14309 | 14309 | 14309 | yes | 1 | 14364 | 14364 | 14364 | no | 1 | 14314 | 14314 | 14314 | yes |
| HAB_2_4a | 1 | 14941 | 14941 | 14941 | no | 1 | 13888 | 13888 | 13888 | no | 1 | 14068 | 14068 | 14068 | no | 1 | 14484 | 14484 | 14484 | no |
| HAB_2_6b | 1 | 14676 | 14676 | 14676 | no | 1 | 14095 | 14095 | 14095 | no | 1 | 13829 | 13829 | 13829 | yes | 1 | 14332 | 14332 | 14332 | yes |
| KLA4_10b | 1 | 14236 | 14236 | 14236 | yes | 1 | 14570 | 14570 | 14570 | no | 1 | 13991 | 13991 | 13991 | no | 1 | 14242 | 14242 | 14242 | yes |
| KLA4_5a | 1 | 14113 | 14113 | 14113 | no | 1 | 14649 | 14649 | 14649 | no | 1 | 14397 | 14397 | 14397 | no | 1 | 14245 | 14245 | 14245 | yes |
| KLA4_9a | 1 | 14492 | 14492 | 14492 | yes | 1 | 14295 | 14295 | 14295 | yes | 1 | 14429 | 14429 | 14429 | no | 1 | 14301 | 14301 | 14301 | yes |
| MUS_6_22a | 1 | 14433 | 14433 | 14433 | no | 1 | 13918 | 13918 | 13918 | no | 1 | 14220 | 14220 | 14220 | no | 1 | 14153 | 14153 | 14153 | no |
| PHU7 | 1 | 14294 | 14294 | 14294 | yes | 1 | 14179 | 14179 | 14179 | no | 1 | 14452 | 14452 | 14452 | yes | 1 | 14520 | 14520 | 14520 | no |
| SRR31631925 | 1 | 14471 | 14471 | 14471 | no | 1 | 14002 | 14002 | 14002 | no | 1 | 13939 | 13939 | 13939 | no | 1 | 14466 | 14466 | 14466 | no |
| SRR31675196 | 1 | 14460 | 14460 | 14460 | yes | 1 | 14020 | 14020 | 14020 | no | 1 | 13998 | 13998 | 13998 | no | 1 | 14560 | 14560 | 14560 | no |
| SRR31675197 | 1 | 14426 | 14426 | 14426 | no | 1 | 14010 | 14010 | 14010 | no | 1 | 13929 | 13929 | 13929 | no | 1 | 14564 | 14564 | 14564 | no |
| SRR31675198 | 1 | 14591 | 14591 | 14591 | no | 1 | 14369 | 14369 | 14369 | no | 1 | 14225 | 14225 | 14225 | no | 1 | 14321 | 14321 | 14321 | yes |
| SRR31675199 | 1 | 14089 | 14089 | 14089 | no | 1 | 14337 | 14337 | 14337 | no | 1 | 13859 | 13859 | 13859 | yes | 1 | 14869 | 14869 | 14869 | no |
| SRR31675200 | 1 | 14579 | 14579 | 14579 | no | 1 | 14073 | 14073 | 14073 | no | 1 | 13989 | 13989 | 13989 | no | 1 | 14627 | 14627 | 14627 | no |
| SRR31675201 | 1 | 14713 | 14713 | 14713 | no | 1 | 14517 | 14517 | 14517 | no | 1 | 13942 | 13942 | 13942 | yes | 1 | 14613 | 14613 | 14613 | no |
| SRR31675202 | 1 | 14499 | 14499 | 14499 | no | 1 | 14002 | 14002 | 14002 | no | 1 | 14114 | 14114 | 14114 | no | 1 | 14332 | 14332 | 14332 | yes |
| SRR31675203 | 1 | 14302 | 14302 | 14302 | yes | 1 | 13656 | 13656 | 13656 | no | 1 | 11912 | 11912 | 11912 | no | 2 | 13637 | 19851 | 13637 | no |
| SRR31675204 | 1 | 14432 | 14432 | 14432 | no | 1 | 13986 | 13986 | 13986 | no | 1 | 13984 | 13984 | 13984 | no | 1 | 14696 | 14696 | 14696 | no |
| SRR31692059 | 4 | 10289 | 17658 | 10289 | no | 3 | 10649 | 15679 | 10649 | no | 3 | 23942 | 29400 | 23942 | no | 2 | 13124 | 16050 | 13124 | no |
| SRR31692061 | 1 | 14757 | 14757 | 14757 | no | 1 | 14390 | 14390 | 14390 | no | 1 | 14318 | 14318 | 14318 | no | 1 | 15092 | 15092 | 15092 | no |
| SRR31692062 | 1 | 14324 | 14324 | 14324 | yes | 1 | 13837 | 13837 | 13837 | yes | 1 | 13860 | 13860 | 13860 | no | 1 | 14330 | 14330 | 14330 | yes |
| SRR31692063 | 1 | 14204 | 14204 | 14204 | no | 1 | 14081 | 14081 | 14081 | no | 1 | 14025 | 14025 | 14025 | no | 1 | 14613 | 14613 | 14613 | no |
| SRR31692064 | 1 | 14566 | 14566 | 14566 | no | 1 | 13959 | 13959 | 13959 | no | 1 | 14031 | 14031 | 14031 | no | 1 | 14487 | 14487 | 14487 | yes |
| SRR31692065 | 1 | 14460 | 14460 | 14460 | yes | 1 | 14008 | 14008 | 14008 | no | 1 | 14061 | 14061 | 14061 | no | 1 | 14460 | 14460 | 14460 | yes |
| SRR31692066 | N/A | N/A | N/A | N/A | N/A | 1 | 14493 | 14493 | 14493 | no | 1 | 14124 | 14124 | 14124 | no | N/A | N/A | N/A | N/A | N/A |
| SRR31692067 | N/A | N/A | N/A | N/A | N/A | 1 | 14292 | 14292 | 14292 | no | 1 | 13981 | 13981 | 13981 | no | 1 | 14471 | 14471 | 14471 | no |
| SRR31692068 | 1 | 14485 | 14485 | 14485 | yes | 1 | 14110 | 14110 | 14110 | no | 1 | 14196 | 14196 | 14196 | no | 1 | 14023 | 14023 | 14023 | no |
| SRR31692069 | 1 | 14723 | 14723 | 14723 | no | 1 | 14226 | 14226 | 14226 | no | 1 | 13942 | 13942 | 13942 | no | 1 | 14469 | 14469 | 14469 | no |
| SRR31692070 | 1 | 14503 | 14503 | 14503 | no | 1 | 13995 | 13995 | 13995 | no | 1 | 13950 | 13950 | 13950 | yes | 1 | 14452 | 14452 | 14452 | yes |
| SRR31692071 | 1 | 5610 | 5610 | 5610 | no | 1 | 14440 | 14440 | 14440 | no | 1 | 14009 | 14009 | 14009 | no | 1 | 14460 | 14460 | 14460 | yes |
| SRR31692072 | 1 | 14840 | 14840 | 14840 | no | 1 | 14504 | 14504 | 14504 | no | 1 | 14111 | 14111 | 14111 | no | 1 | 14498 | 14498 | 14498 | no |
| SRR31692073 | 1 | 14530 | 14530 | 14530 | no | 1 | 14253 | 14253 | 14253 | no | 1 | 14343 | 14343 | 14343 | yes | N/A | N/A | N/A | N/A | N/A |
| SRR31692074 | 1 | 14680 | 14680 | 14680 | no | 1 | 13827 | 13827 | 13827 | yes | 1 | 13859 | 13859 | 13859 | no | 1 | 14326 | 14326 | 14326 | yes |
| SRR31692075 | 1 | 14460 | 14460 | 14460 | yes | 3 | 13624 | 19708 | 13624 | no | 3 | 13648 | 19732 | 13648 | no | 1 | 27134 | 27134 | 27134 | no |
| SRR31692076 | 1 | 14520 | 14520 | 14520 | yes | 1 | 13968 | 13968 | 13968 | no | 1 | 13955 | 13955 | 13955 | no | 1 | 14418 | 14418 | 14418 | yes |
| SRR31692077 | 1 | 14284 | 14284 | 14284 | no | 1 | 14077 | 14077 | 14077 | no | 1 | 14164 | 14164 | 14164 | no | 1 | 14385 | 14385 | 14385 | no |
| SRR31692078 | 1 | 14464 | 14464 | 14464 | no | 1 | 14217 | 14217 | 14217 | no | 1 | 13992 | 13992 | 13992 | no | 1 | 14436 | 14436 | 14436 | yes |
| SRR31757516 | 1 | 14302 | 14302 | 14302 | yes | 1 | 14048 | 14048 | 14048 | no | 1 | 13868 | 13868 | 13868 | no | 1 | 14298 | 14298 | 14298 | yes |
| SRR31757517 | 1 | 14554 | 14554 | 14554 | no | 1 | 14465 | 14465 | 14465 | no | 1 | 14030 | 14030 | 14030 | no | 1 | 14454 | 14454 | 14454 | yes |
| SRR31757518 | 1 | 14643 | 14643 | 14643 | no | 1 | 13951 | 13951 | 13951 | yes | 1 | 13996 | 13996 | 13996 | no | 1 | 14443 | 14443 | 14443 | yes |
| SRR31757519 | 1 | 13990 | 13990 | 13990 | no | 1 | 13817 | 13817 | 13817 | no | 1 | 13881 | 13881 | 13881 | yes | 1 | 14676 | 14676 | 14676 | no |
| SRR31757520 | 1 | 14484 | 14484 | 14484 | no | 1 | 13991 | 13991 | 13991 | no | 1 | 13859 | 13859 | 13859 | yes | 1 | 14304 | 14304 | 14304 | no |
| SRR31757521 | 1 | 14858 | 14858 | 14858 | no | 1 | 13962 | 13962 | 13962 | no | 1 | 14279 | 14279 | 14279 | no | 1 | 14466 | 14466 | 14466 | yes |
| SRR31757522 | 1 | 14224 | 14224 | 14224 | no | 1 | 13968 | 13968 | 13968 | no | 1 | 13879 | 13879 | 13879 | yes | 1 | 14448 | 14448 | 14448 | yes |
| SRR31757523 | 1 | 14628 | 14628 | 14628 | no | 1 | 13989 | 13989 | 13989 | no | 1 | 13966 | 13966 | 13966 | yes | 1 | 14692 | 14692 | 14692 | no |
| SRR31757524 | 1 | 14660 | 14660 | 14660 | no | 1 | 14096 | 14096 | 14096 | no | 1 | 13958 | 13958 | 13958 | yes | 1 | 14581 | 14581 | 14581 | no |
| SRR31757525 | 1 | 14068 | 14068 | 14068 | no | 1 | 13967 | 13967 | 13967 | no | 1 | 13936 | 13936 | 13936 | yes | 1 | 14144 | 14144 | 14144 | no |
| SRR31757526 | 1 | 14572 | 14572 | 14572 | no | 1 | 13929 | 13929 | 13929 | no | 1 | 13936 | 13936 | 13936 | yes | 1 | 14616 | 14616 | 14616 | no |
| SRR31757527 | 1 | 14051 | 14051 | 14051 | no | 1 | 14056 | 14056 | 14056 | no | 1 | 13860 | 13860 | 13860 | no | 1 | 14331 | 14331 | 14331 | yes |
| SRR31757528 | 1 | 14628 | 14628 | 14628 | no | 1 | 13999 | 13999 | 13999 | no | 1 | 13988 | 13988 | 13988 | no | 1 | 14835 | 14835 | 14835 | yes |
| SRR31757529 | 1 | 14514 | 14514 | 14514 | no | 1 | 14030 | 14030 | 14030 | no | 1 | 14037 | 14037 | 14037 | no | 1 | 14660 | 14660 | 14660 | no |
| SRR31757530 | 1 | 14110 | 14110 | 14110 | no | 1 | 13990 | 13990 | 13990 | no | 1 | 14020 | 14020 | 14020 | no | 1 | 14066 | 14066 | 14066 | no |
| SRR31757531 | 1 | 14105 | 14105 | 14105 | no | 1 | 13996 | 13996 | 13996 | no | 1 | 13981 | 13981 | 13981 | no | 1 | 14105 | 14105 | 14105 | no |
| SRR31757532 | 1 | 14213 | 14213 | 14213 | no | 1 | 13985 | 13985 | 13985 | no | 1 | 14015 | 14015 | 14015 | no | 1 | 14193 | 14193 | 14193 | no |
| SRR31757533 | 1 | 14933 | 14933 | 14933 | no | 1 | 13997 | 13997 | 13997 | no | 1 | 14028 | 14028 | 14028 | no | 1 | 14217 | 14217 | 14217 | no |
| SRR31757534 | 1 | 14017 | 14017 | 14017 | no | 1 | 13957 | 13957 | 13957 | no | 1 | 13949 | 13949 | 13949 | yes | 1 | 13963 | 13963 | 13963 | no |
| SRR31757535 | 1 | 14047 | 14047 | 14047 | no | 1 | 13987 | 13987 | 13987 | no | 1 | 13890 | 13890 | 13890 | no | 1 | 14508 | 14508 | 14508 | no |
| SRR31757536 | 1 | 14615 | 14615 | 14615 | no | 1 | 14093 | 14093 | 14093 | no | 1 | 14004 | 14004 | 14004 | no | 1 | 14133 | 14133 | 14133 | no |
| SRR31757537 | 1 | 14454 | 14454 | 14454 | yes | 1 | 13980 | 13980 | 13980 | no | 1 | 13997 | 13997 | 13997 | no | 1 | 14470 | 14470 | 14470 | yes |
| SRR31757538 | 1 | 14131 | 14131 | 14131 | no | 1 | 13986 | 13986 | 13986 | no | 1 | 14230 | 14230 | 14230 | no | 1 | 14796 | 14796 | 14796 | no |
| SRR31757539 | 1 | 14491 | 14491 | 14491 | yes | 1 | 14156 | 14156 | 14156 | no | 1 | 13980 | 13980 | 13980 | no | 1 | 14586 | 14586 | 14586 | no |
| SRR31757540 | 1 | 14508 | 14508 | 14508 | yes | 1 | 14057 | 14057 | 14057 | no | 1 | 14029 | 14029 | 14029 | no | 1 | 14502 | 14502 | 14502 | yes |
| SRR31757541 | 1 | 14058 | 14058 | 14058 | no | 1 | 13948 | 13948 | 13948 | yes | 1 | 13979 | 13979 | 13979 | no | 1 | 15016 | 15016 | 15016 | no |
| SRR31757542 | 1 | 14519 | 14519 | 14519 | yes | 1 | 14284 | 14284 | 14284 | no | 1 | 13944 | 13944 | 13944 | no | 1 | 14713 | 14713 | 14713 | no |
| SRR31757543 | 1 | 14007 | 14007 | 14007 | no | 1 | 14023 | 14023 | 14023 | no | 1 | 14047 | 14047 | 14047 | no | 1 | 14715 | 14715 | 14715 | no |
| SRR31757544 | 1 | 14601 | 14601 | 14601 | no | 1 | 14110 | 14110 | 14110 | no | 1 | 13865 | 13865 | 13865 | yes | 1 | 14386 | 14386 | 14386 | no |
| SRR31757545 | 1 | 14515 | 14515 | 14515 | no | 1 | 14015 | 14015 | 14015 | no | 1 | 14386 | 14386 | 14386 | no | 1 | 22628 | 22628 | 22628 | no |

| | | | | | | | | | | | | | | | | | | | | |
|---|---|---|---|---|---|---|---|---|---|---|---|---|---|---|---|---|---|---|---|---|
| SRR31757546 | 1 | 14334 | 14334 | 14334 | yes | 1 | 14334 | 14334 | 14334 | yes | 1 | 13943 | 13943 | 13943 | no | 1 | 14136 | 14136 | 14136 | no |
| SRR31757547 | 1 | 14104 | 14104 | 14104 | no | 1 | 14114 | 14114 | 14114 | no | 1 | 14005 | 14005 | 14005 | no | 1 | 14684 | 14684 | 14684 | no |
| SRR8419293 | 1 | 14280 | 14280 | 14280 | yes | 1 | 14353 | 14353 | 14353 | no | 1 | 14212 | 14212 | 14212 | no | 1 | 14278 | 14278 | 14278 | yes |
| SRR8419294 | 1 | 14312 | 14312 | 14312 | yes | 1 | 14154 | 14154 | 14154 | yes | 1 | 14221 | 14221 | 14221 | no | 1 | 14310 | 14310 | 14310 | yes |
| SRR8419295 | 1 | 14364 | 14364 | 14364 | yes | 1 | 14043 | 14043 | 14043 | no | 1 | 13890 | 13890 | 13890 | no | 1 | 14390 | 14390 | 14390 | no |
| SRR8419296 | 1 | 14378 | 14378 | 14378 | no | 1 | 13911 | 13911 | 13911 | no | 1 | 14112 | 14112 | 14112 | yes | 1 | 14399 | 14399 | 14399 | no |
| SRR8419297 | 1 | 14315 | 14315 | 14315 | yes | 1 | 14275 | 14275 | 14275 | no | 1 | 13908 | 13908 | 13908 | no | 1 | 14321 | 14321 | 14321 | yes |
| SRR8419298 | 1 | 14348 | 14348 | 14348 | yes | 1 | 14017 | 14017 | 14017 | no | 1 | 13955 | 13955 | 13955 | no | 1 | 14344 | 14344 | 14344 | yes |
| SRR8419299 | 1 | 14089 | 14089 | 14089 | no | 1 | 13890 | 13890 | 13890 | no | 1 | 13853 | 13853 | 13853 | yes | 1 | 14276 | 14276 | 14276 | yes |
| SRR8419300 | 1 | 14397 | 14397 | 14397 | no | 1 | 14372 | 14372 | 14372 | no | 1 | 14010 | 14010 | 14010 | yes | 1 | 14395 | 14395 | 14395 | no |
| SRR8419301 | 1 | 13931 | 13931 | 13931 | no | 1 | 14049 | 14049 | 14049 | no | 1 | 14264 | 14264 | 14264 | no | 1 | 14365 | 14365 | 14365 | no |
| SRR8419302 | 1 | 14388 | 14388 | 14388 | no | 1 | 14332 | 14332 | 14332 | yes | 1 | 14004 | 14004 | 14004 | yes | 1 | 14340 | 14340 | 14340 | yes |
| SRR8419303 | 1 | 14334 | 14334 | 14334 | yes | 1 | 13894 | 13894 | 13894 | no | 1 | 13892 | 13892 | 13892 | no | 1 | 14356 | 14356 | 14356 | no |
| SRR8419304 | 1 | 14420 | 14420 | 14420 | no | 1 | 14382 | 14382 | 14382 | no | 1 | 13856 | 13856 | 13856 | yes | 1 | 13900 | 13900 | 13900 | no |
| SRR8419305 | 1 | 14388 | 14388 | 14388 | no | 1 | 13940 | 13940 | 13940 | no | 1 | 13911 | 13911 | 13911 | no | 1 | 14423 | 14423 | 14423 | no |
| SRR8419306 | 1 | 14354 | 14354 | 14354 | no | 1 | 13923 | 13923 | 13923 | no | 1 | 14082 | 14082 | 14082 | no | 1 | 14352 | 14352 | 14352 | yes |
| SRR8419307 | 1 | 14407 | 14407 | 14407 | no | 1 | 14060 | 14060 | 14060 | no | 1 | 13882 | 13882 | 13882 | no | 1 | 14471 | 14471 | 14471 | no |
| SRR8419308 | 1 | 14389 | 14389 | 14389 | no | 1 | 14328 | 14328 | 14328 | no | 1 | 13856 | 13856 | 13856 | yes | 1 | 14365 | 14365 | 14365 | yes |
| SRR8419309 | 1 | 14445 | 14445 | 14445 | no | 1 | 14382 | 14382 | 14382 | no | 1 | 13892 | 13892 | 13892 | no | 1 | 14452 | 14452 | 14452 | no |
| SRR8419310 | 1 | 14400 | 14400 | 14400 | no | 1 | 14446 | 14446 | 14446 | no | 1 | 13851 | 13851 | 13851 | yes | 1 | 14360 | 14360 | 14360 | yes |
| SRR8419311 | 1 | 14371 | 14371 | 14371 | no | 1 | 13901 | 13901 | 13901 | no | 1 | 13851 | 13851 | 13851 | yes | 1 | 13969 | 13969 | 13969 | no |
| SRR8419312 | 1 | 14314 | 14314 | 14314 | yes | 1 | 14328 | 14328 | 14328 | no | 1 | 14133 | 14133 | 14133 | no | 1 | 14310 | 14310 | 14310 | yes |
| SRR8419313 | 1 | 14335 | 14335 | 14335 | yes | 1 | 14074 | 14074 | 14074 | no | 1 | 13852 | 13852 | 13852 | yes | 1 | 14327 | 14327 | 14327 | yes |
| SRR8419314 | 1 | 14727 | 14727 | 14727 | no | 1 | 13890 | 13890 | 13890 | yes | 1 | 14221 | 14221 | 14221 | no | 1 | 14450 | 14450 | 14450 | no |
| SRR8419315 | 1 | 14418 | 14418 | 14418 | no | 1 | 13961 | 13961 | 13961 | no | 1 | 13853 | 13853 | 13853 | yes | 1 | 14340 | 14340 | 14340 | yes |
| SRR8419316 | 1 | 14358 | 14358 | 14358 | no | 1 | 14444 | 14444 | 14444 | no | 1 | 13853 | 13853 | 13853 | yes | 1 | 14344 | 14344 | 14344 | yes |
| SRR8419317 | 1 | 14340 | 14340 | 14340 | no | 1 | 14356 | 14356 | 14356 | no | 1 | 14106 | 14106 | 14106 | yes | 1 | 14387 | 14387 | 14387 | no |
| SRR8419318 | 1 | 14480 | 14480 | 14480 | no | 1 | 13897 | 13897 | 13897 | no | 1 | 13852 | 13852 | 13852 | yes | 1 | 13892 | 13892 | 13892 | yes |
| SRR8419319 | 1 | 14364 | 14364 | 14364 | no | 1 | 14153 | 14153 | 14153 | no | 1 | 13847 | 13847 | 13847 | no | 1 | 14336 | 14336 | 14336 | yes |
| SRR8419320 | 1 | 14369 | 14369 | 14369 | no | 1 | 13923 | 13923 | 13923 | no | 1 | 13851 | 13851 | 13851 | yes | 1 | 13983 | 13983 | 13983 | no |
| SRR8419321 | 1 | 14358 | 14358 | 14358 | yes | 1 | 14285 | 14285 | 14285 | no | 1 | 13935 | 13935 | 13935 | yes | 1 | 14352 | 14352 | 14352 | yes |
| SRR8419322 | 1 | 13933 | 13933 | 13933 | no | 1 | 13973 | 13973 | 13973 | no | 1 | 13895 | 13895 | 13895 | no | 1 | 13933 | 13933 | 13933 | no |
| SRR8419324 | 1 | 14329 | 14329 | 14329 | yes | 1 | 13931 | 13931 | 13931 | no | 1 | 14004 | 14004 | 14004 | no | 1 | 13931 | 13931 | 13931 | no |
| SRR8419326 | 1 | 13889 | 13889 | 13889 | yes | 1 | 14360 | 14360 | 14360 | no | 1 | 14028 | 14028 | 14028 | yes | 1 | 14372 | 14372 | 14372 | yes |
| SRR8419327 | 1 | 14470 | 14470 | 14470 | no | 1 | 14332 | 14332 | 14332 | yes | 1 | 13930 | 13930 | 13930 | no | 1 | 14384 | 14384 | 14384 | yes |
| SRR8419328 | 1 | 14273 | 14273 | 14273 | no | 1 | 14413 | 14413 | 14413 | no | 1 | 14112 | 14112 | 14112 | yes | 1 | 14412 | 14412 | 14412 | no |
| SRR8419329 | 1 | 14283 | 14283 | 14283 | no | 1 | 14119 | 14119 | 14119 | no | 1 | 14155 | 14155 | 14155 | no | 1 | 14426 | 14426 | 14426 | no |
| SRR8419330 | 1 | 13965 | 13965 | 13965 | no | 1 | 13964 | 13964 | 13964 | no | 1 | 14052 | 14052 | 14052 | no | 1 | 14318 | 14318 | 14318 | yes |
| SRR8419331 | 1 | 14392 | 14392 | 14392 | no | 1 | 14243 | 14243 | 14243 | no | 1 | 14023 | 14023 | 14023 | yes | 1 | 14351 | 14351 | 14351 | no |
| SRR8419332 | 1 | 13924 | 13924 | 13924 | no | 1 | 14345 | 14345 | 14345 | no | 1 | 14039 | 14039 | 14039 | no | 1 | 14340 | 14340 | 14340 | no |
| SRR8419333 | 1 | 14398 | 14398 | 14398 | no | 1 | 14431 | 14431 | 14431 | no | 1 | 13889 | 13889 | 13889 | no | 1 | 14329 | 14329 | 14329 | yes |
| SRR8419334 | 1 | 14266 | 14266 | 14266 | yes | 1 | 14387 | 14387 | 14387 | no | 1 | 14120 | 14120 | 14120 | no | 1 | 14360 | 14360 | 14360 | yes |
| SRR8419335 | 1 | 13865 | 13865 | 13865 | yes | 1 | 14306 | 14306 | 14306 | yes | 1 | 13861 | 13861 | 13861 | yes | 1 | 14394 | 14394 | 14394 | no |
| SRR8419336 | 1 | 14422 | 14422 | 14422 | no | 1 | 14320 | 14320 | 14320 | no | 1 | 14273 | 14273 | 14273 | no | 1 | 14313 | 14313 | 14313 | yes |
| SRR8419338 | 1 | 14360 | 14360 | 14360 | yes | 1 | 14385 | 14385 | 14385 | no | 1 | 14038 | 14038 | 14038 | no | 1 | 14322 | 14322 | 14322 | yes |
| SRR8419339 | 1 | 14274 | 14274 | 14274 | no | 1 | 14210 | 14210 | 14210 | yes | 1 | 13969 | 13969 | 13969 | no | 1 | 14386 | 14386 | 14386 | no |
| SRR8419340 | 1 | 13903 | 13903 | 13903 | no | 1 | 13885 | 13885 | 13885 | no | 1 | 13852 | 13852 | 13852 | yes | 1 | 14347 | 14347 | 14347 | yes |
| SRR8419341 | 1 | 14333 | 14333 | 14333 | yes | 1 | 14457 | 14457 | 14457 | no | 1 | 13996 | 13996 | 13996 | yes | 1 | 14333 | 14333 | 14333 | yes |
| SRR8419342 | 1 | 13956 | 13956 | 13956 | no | 1 | 14395 | 14395 | 14395 | no | 1 | 13915 | 13915 | 13915 | no | 1 | 14068 | 14068 | 14068 | no |
| SRR8419343 | 1 | 13175 | 13175 | 13175 | no | 1 | 13993 | 13993 | 13993 | no | 1 | 14101 | 14101 | 14101 | no | 1 | 13889 | 13889 | 13889 | yes |
| SRR8419344 | 1 | 14310 | 14310 | 14310 | yes | 1 | 14409 | 14409 | 14409 | no | 1 | 14102 | 14102 | 14102 | yes | 1 | 14310 | 14310 | 14310 | yes |
| SRR8419345 | 1 | 14389 | 14389 | 14389 | yes | 1 | 13962 | 13962 | 13962 | no | 1 | 13868 | 13868 | 13868 | yes | 1 | 14452 | 14452 | 14452 | no |
| SRR8419347 | 1 | 13931 | 13931 | 13931 | no | 1 | 14347 | 14347 | 14347 | no | 1 | 14074 | 14074 | 14074 | no | 1 | 14430 | 14430 | 14430 | no |
| SRR8419348 | N/A | N/A | N/A | N/A | N/A | 1 | 13878 | 13878 | 13878 | yes | 1 | 13856 | 13856 | 13856 | yes | 1 | 13932 | 13932 | 13932 | no |
| SRR8419349 | 1 | 13880 | 13880 | 13880 | yes | 1 | 12702 | 12702 | 12702 | no | 1 | 13830 | 13830 | 13830 | yes | 1 | 14365 | 14365 | 14365 | yes |
| SRR8419350 | 1 | 14363 | 14363 | 14363 | no | 1 | 13881 | 13881 | 13881 | yes | 1 | 14112 | 14112 | 14112 | yes | 1 | 14338 | 14338 | 14338 | no |
| SRR8419351 | 1 | 14330 | 14330 | 14330 | no | 1 | 14336 | 14336 | 14336 | yes | 1 | 14149 | 14149 | 14149 | no | 1 | 14270 | 14270 | 14270 | yes |
| SRR8419352 | 1 | 14334 | 14334 | 14334 | yes | 1 | 13851 | 13851 | 13851 | yes | 1 | 13980 | 13980 | 13980 | no | 1 | 14364 | 14364 | 14364 | no |
| SRR8419353 | 1 | 13911 | 13911 | 13911 | no | 1 | 14364 | 14364 | 14364 | no | 1 | 13855 | 13855 | 13855 | yes | 1 | 14415 | 14415 | 14415 | no |
| SRR8419354 | 1 | 13891 | 13891 | 13891 | no | 1 | 14002 | 14002 | 14002 | no | 1 | 13853 | 13853 | 13853 | yes | 1 | 14491 | 14491 | 14491 | no |
| SRR8419355 | 1 | 13917 | 13917 | 13917 | no | 1 | 14301 | 14301 | 14301 | no | 1 | 14333 | 14333 | 14333 | no | 1 | 14370 | 14370 | 14370 | no |
| SRR8419356 | 1 | 14118 | 14118 | 14118 | yes | 1 | 14364 | 14364 | 14364 | yes | 1 | 14147 | 14147 | 14147 | no | 1 | 14120 | 14120 | 14120 | yes |
| SRR8419357 | 1 | 14417 | 14417 | 14417 | no | 1 | 14389 | 14389 | 14389 | no | 1 | 13851 | 13851 | 13851 | yes | 1 | 14348 | 14348 | 14348 | yes |
| SRR8419358 | 1 | 14383 | 14383 | 14383 | no | 1 | 14461 | 14461 | 14461 | no | 1 | 13915 | 13915 | 13915 | no | 1 | 14322 | 14322 | 14322 | yes |
| SRR8419359 | 1 | 14381 | 14381 | 14381 | no | 1 | 13902 | 13902 | 13902 | no | 1 | 13889 | 13889 | 13889 | no | 1 | 14371 | 14371 | 14371 | no |
| SRR8419360 | 1 | 14340 | 14340 | 14340 | yes | 1 | 13845 | 13845 | 13845 | yes | 1 | 13979 | 13979 | 13979 | yes | 1 | 13942 | 13942 | 13942 | no |
| TrentoS7 | 1 | 14288 | 14288 | 14288 | yes | 1 | 14383 | 14383 | 14383 | no | 1 | 13914 | 13914 | 13914 | no | 1 | 14288 | 14288 | 14288 | yes |
| TrentoS8 | 1 | 14286 | 14286 | 14286 | yes | 1 | 13861 | 13861 | 13861 | no | 1 | 13852 | 13852 | 13852 | yes | 1 | 14487 | 14487 | 14487 | no |
| UK16F13605_b | 1 | 14649 | 14649 | 14649 | no | 1 | 14418 | 14418 | 14418 | no | 1 | 13968 | 13968 | 13968 | no | 1 | 14357 | 14357 | 14357 | yes |
| circular | 43/146 | | | | | 22/149 | | | | | 48/149 | | | | | 67/147 | | | | |

failed

no full mtDNA

multiple contigs including complete mtDNA contig

**Supplementary Table S3: Summary of mapping metrics for *De Novo* assembled mitochondrial genomes**.
Analysis of subsampled sequencing reads mapped back to their respective assemblies using Bowtie2 *successfully covering the mitochondrial genome

| Sample ID | Length | Number of reads* | Coverage | Mean depth per base | Mean base quality | Mean mapping quality |
|---|---|---|---|---|---|---|
| **AL12SLB** | 14107 | 2664836 | 99.84 | 28322.1 | 35.4 | 41.7 |
| **AL17SLB** | 14314 | 3805662 | 100 | 39858.1 | 35.6 | 41.7 |
| **AL2SLB** | 14266 | 1793415 | 100 | 18844.3 | 35.4 | 41.7 |
| **Bunawan S10** | 14300 | 5344565 | 100 | 54755.6 | 35.1 | 41.6 |
| **Bunawan S9** | 14484 | 772984 | 100 | 7741.83 | 35 | 41.4 |
| **DE H1a** | 14429 | 1298423 | 99.99 | 13465 | 39.5 | 41.6 |
| **DE P11** | 14089 | 3596447 | 100 | 37962.4 | 39.5 | 41.8 |
| **DE P14b** | 14027 | 8896601 | 100 | 94984.1 | 39.5 | 41.8 |
| **DE P17a** | 13833 | 3810451 | 100 | 40958 | 39.4 | 41.7 |
| **DE P22** | 14014 | 6133917 | 100 | 65531.1 | 39.5 | 41.8 |
| **DE P24** | 13967 | 8390230 | 100 | 89825.7 | 39.5 | 41.8 |
| **DE P6** | 14082 | 4837586 | 100 | 51326.7 | 39.5 | 41.8 |
| **DE P9** | 14418 | 6110136 | 100 | 63401.8 | 39.5 | 41.7 |
| **HAB 2.3b** | 14300 | 5612739 | 100 | 58197.3 | 35.6 | 41.6 |
| **HAB 2.4a** | 13882 | 5389733 | 100 | 57195.5 | 35.6 | 41.6 |
| **HAB 2.6b** | 13968 | 1676488 | 100 | 17803.3 | 35.6 | 41.7 |
| **KLA 4.10b** | 14236 | 6550649 | 100 | 68626.4 | 35.6 | 41.6 |
| **KLA 4.5a** | 13972 | 4356068 | 100 | 46624.1 | 35.8 | 41.7 |
| **KLA 4.9a** | 14299 | 15101947 | 100 | 158022 | 35.8 | 41.8 |
| **MUS 6.22a** | 13895 | 1606977 | 100 | 17125.8 | 35.7 | 41.6 |
| **PHU7** | 13943 | 2346481 | 100 | 25228 | 35.4 | 41.5 |
| **Trento S7** | 14384 | 379796 | 100 | 3651.48 | 35.1 | 41.4 |
| **Trento S8** | 14282 | 108584 | 100 | 937.03 | 35.2 | 41.4 |
| **SRR31675200** | 14063 | 1974429 | 100 | 21053.6 | 35.6 | 41.7 |
| **SRR31675201** | 14495 | 1456871 | 100 | 15070.9 | 35.6 | 41.7 |
| **SRR31675202** | 14338 | 1646436 | 100 | 17219.3 | 34.8 | 41.4 |
| **SRR8419293** | 14322 | 14078167 | 100 | 92275.4 | 37 | 41.7 |
| **SRR8419294** | 14431 | 5195710 | 100 | 33609.6 | 36.5 | 41.5 |
| **SRR8419295** | 13942 | 7420389 | 100 | 49482.6 | 36.4 | 41.6 |

**Supplementary Table S4: Accession number of ascarid species used for mtDNA enrichment using Deacon**

| Species | Accession Number(s) |
|---|---|
| *Ascaris lumbricoides* | AP017677.1<br>HQ704900.1<br>KY045803.1<br>SRR31757522<br>SRR31757533 |
| *Parascaris equorum* | AP017696.1 |
| *Ascaris sp. chimpanzee-cA* | KC839986.1 |
| *Ascaris suum* | NC_001327<br>HQ704901.1<br>KY045800.1<br>KY045804.1 |
| *Baylisascaris procyonis* | NC_016200 |
| *Toxascaris leonina TLST* | MW560284.1 |

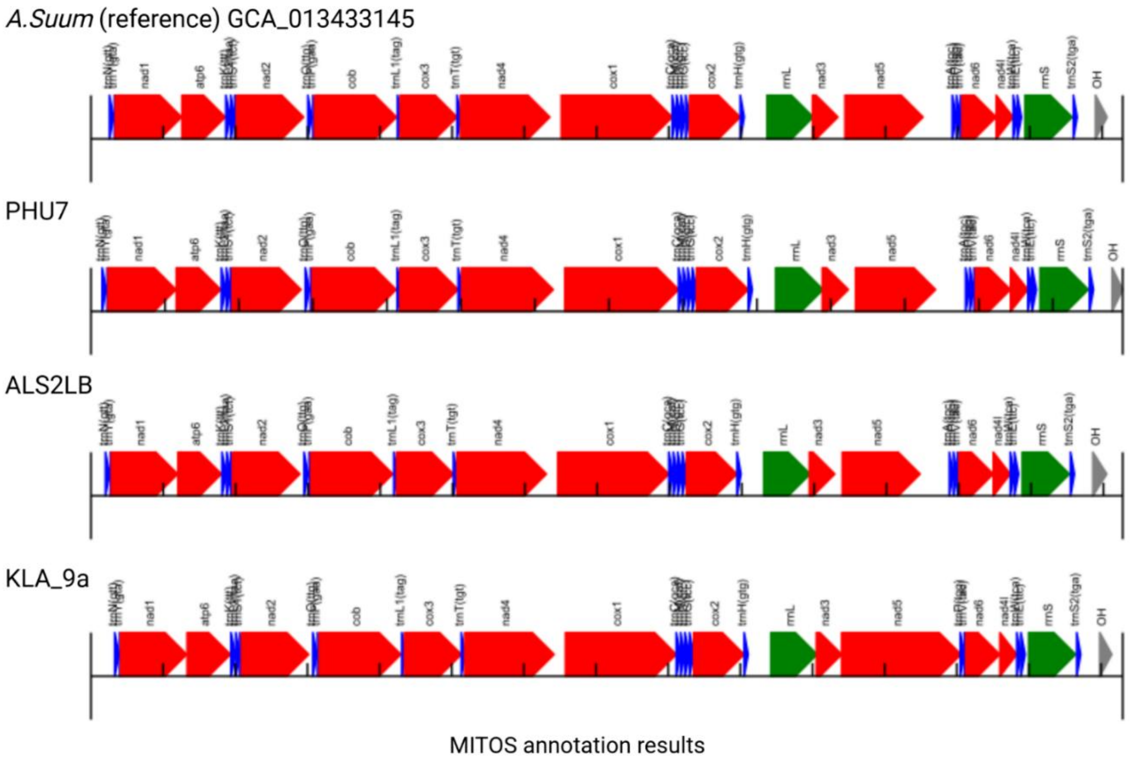


**Supplementary Figure S1**. Assembly of the mitochondrial genome of *Ascaris* spp. generated using MITOZ with annotation from MITOS2. The edited mitochondrial genome was linearised for visualisation, and ordered to start at *nad1* and displays the relative positions and orientations of annotated genes. Protein-coding genes are shown in red, transfer RNA (tRNA) genes in blue, and ribosomal RNA (rRNA) genes in green. The grey region at the right end (labelled "OH") represents the putative AT-rich control region, corresponding to the origin of replication for the heavy strand. Gene annotations and boundaries were predicted using the invertebrate mitochondrial genetic code. Direction of arrows indicates the transcriptional orientation of each gene

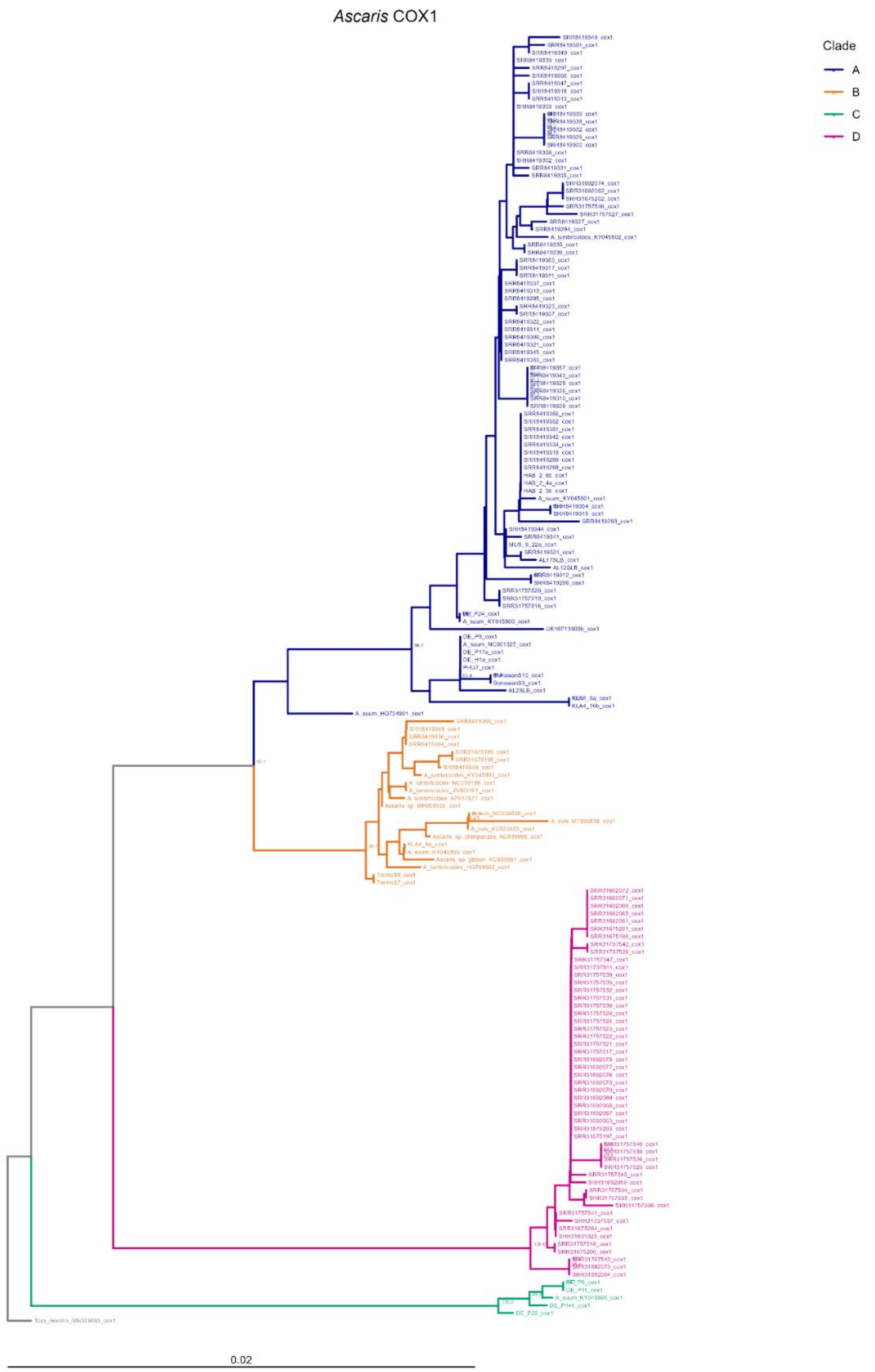


**Supplementary Figure S2:** ***Neighbour-joining tree of cox1 from Ascaris samples*.**
Bootstrap values (>80%) from 1,000 replicates are shown at internal nodes. Tip labels correspond to individual sample identifiers. The tree is drawn to scale, with branch lengths in the same units as those of the evolutionary distances to infer the phylogenomic tree. The evolutionary distances were computed using the Maximum Composite Likelihood Method and are in the units of the number of base substitutions per site. The analysis encompassed 168 sequences aligned using MEGA12. Clades A–D were defined based on phylogenetic clustering. Colour indicates clade assignment as follows: blue (A), orange (B), green (C), and pink (D). The tree is rooted using the outgroup *Toxascaris leonina* MN329693 (scale bar: 0.02 nucleotide substitutions per base).

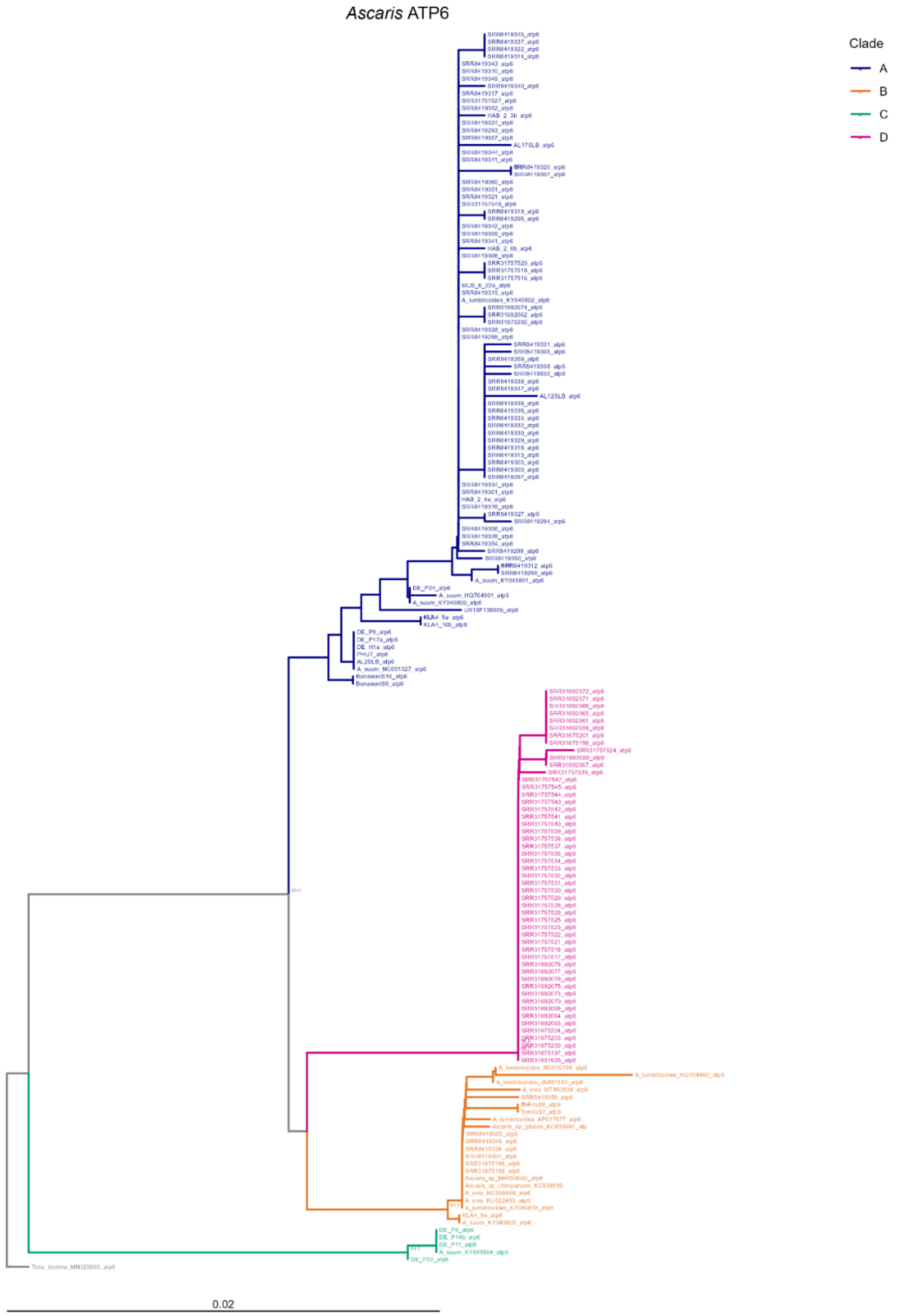


**Supplementary Figure S3** ***Neighbour-joining tree of atp6 from Ascaris samples.***
Bootstrap values (>80%) from 1,000 replicates are shown at internal nodes. Tip labels correspond to individual sample identifiers. The tree is drawn to scale, with branch lengths in the same units as those of the evolutionary distances to infer the phylogenomic tree. The evolutionary distances were computed using the Maximum Composite Likelihood Method and are in the units of the number of base substitutions per site. The analysis encompassed 168 sequences aligned using MEGA12. Clades A–D were defined based on phylogenetic clustering. Colour indicates clade assignment as follows: blue (A), orange (B), green (C), and pink (D). The tree is rooted using the outgroup *Toxascaris leonina* MN329693 (scale bar: 0.02 nucleotide substitutions per base).

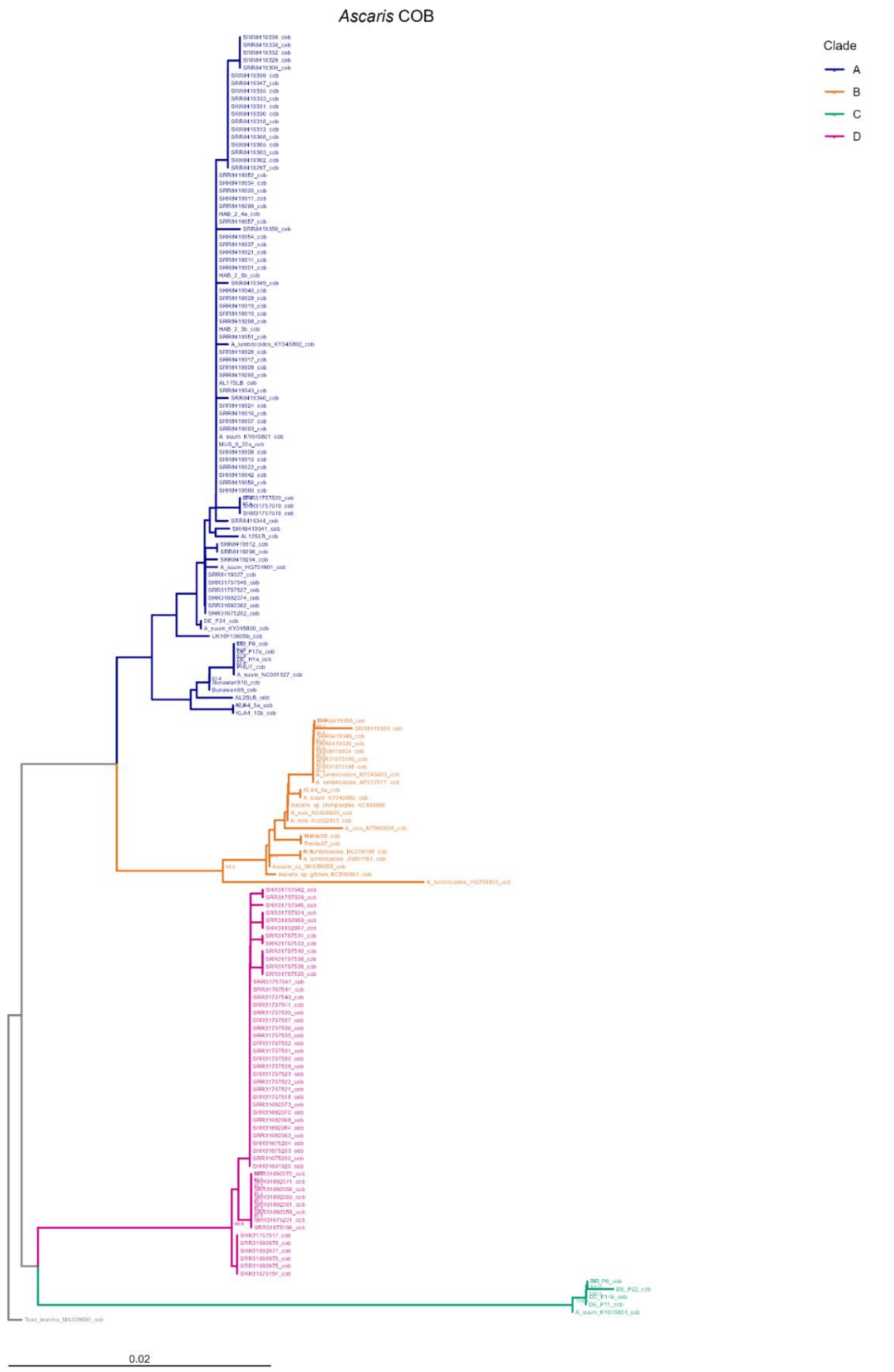


**Supplementary Figure S4** ***Neighbour-joining tree of cob from Ascaris samples.***
Bootstrap values (>80%) from 1,000 replicates are shown at internal nodes. Tip labels correspond to individual sample identifiers. The tree is drawn to scale, with branch lengths in the same units as those of the evolutionary distances to infer the phylogenomic tree. The evolutionary distances were computed using the Maximum Composite Likelihood Method and are in the units of the number of base substitutions per site. The analysis encompassed 168 sequences, aligned using MEGA12. Clades A–D were defined based on phylogenetic clustering. Colour indicates clade assignment as follows: blue (A), orange (B), green (C), and pink (D). The tree is rooted using the outgroup *Toxascaris leonina* MN329693 (scale bar: 0.02 nucleotide substitutions per base).

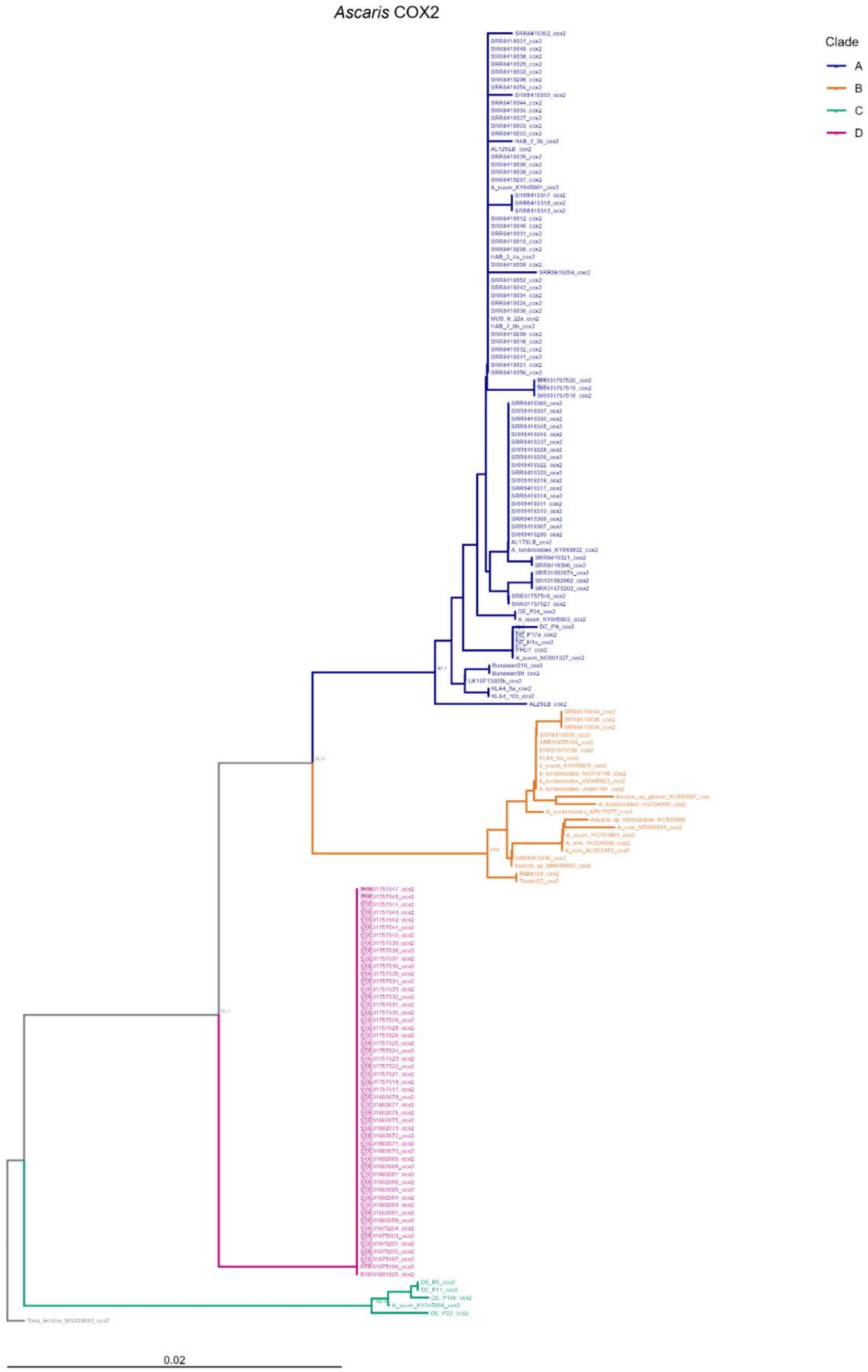


**Supplementary Figure S5** ***Neighbour-joining tree of cox2 from Ascaris samples.***
Bootstrap values (>80%) from 1,000 replicates are shown at internal nodes. Tip labels correspond to individual sample identifiers. The tree is drawn to scale, with branch lengths in the same units as those of the evolutionary distances to infer the phylogenomic tree. The evolutionary distances were computed using the Maximum Composite Likelihood Method and are in the units of the number of base substitutions per site. The analysis encompassed 168 sequences, aligned using MEGA12. Clades A–D were defined based on phylogenetic clustering. Colour indicates clade assignment as follows: blue (A), orange (B), green (C), and pink (D). The tree is rooted using the outgroup *Toxascaris leonina* MN329693 (scale bar: 0.02 nucleotide substitutions per base).

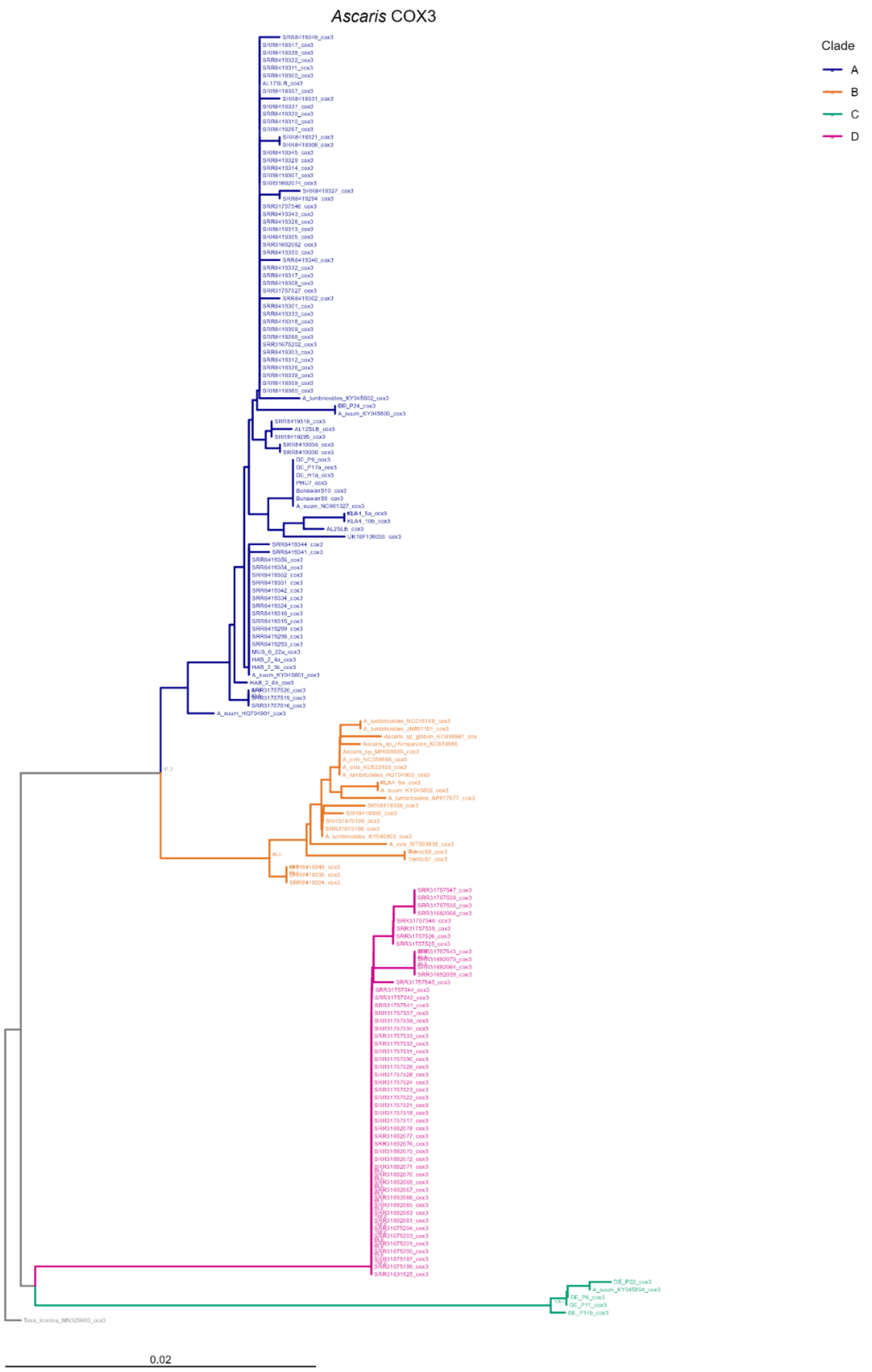


**Supplementary Figure S6** ***Neighbour-joining tree of cox3 from Ascaris samples*.**
Bootstrap values (>80%) from 1,000 replicates are shown at internal nodes. Tip labels correspond to individual sample identifiers. The tree is drawn to scale, with branch lengths in the same units as those of the evolutionary distances to infer the phylogenomic tree. The evolutionary distances were computed using the Maximum Composite Likelihood Method and are in the units of the number of base substitutions per site. The analysis encompassed 168 sequences, aligned using MEGA12. Clades A–D were defined based on phylogenetic clustering. Colour indicates clade assignment as follows: blue (A), orange (B), green (C), and pink (D). The tree is rooted using the outgroup *Toxascaris leonina* MN329693 (scale bar: 0.02 nucleotide substitutions per base).

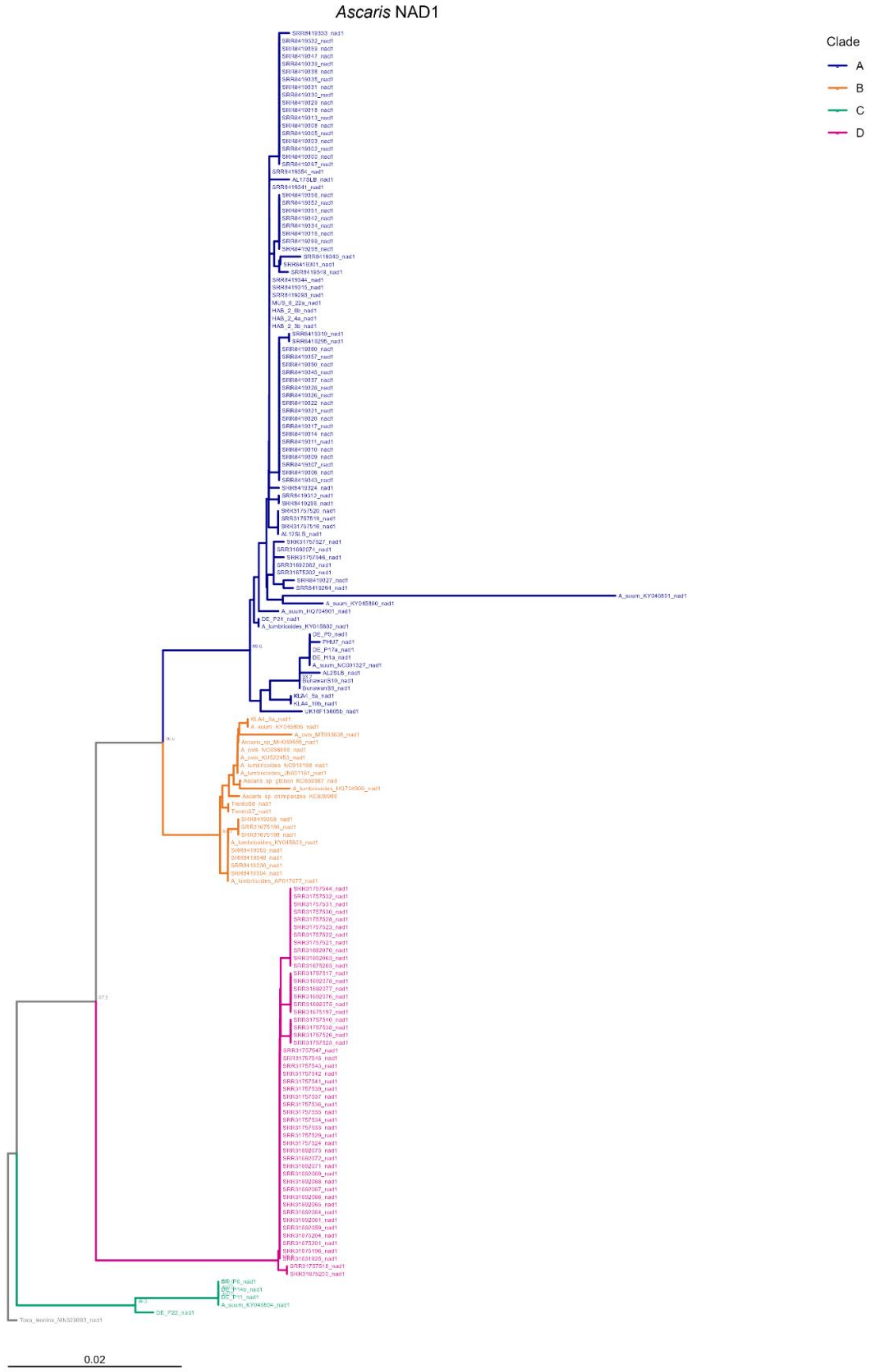


**Supplementary Figure S7** ***Neighbour-joining tree of nad1 from Ascaris samples.***
Bootstrap values (>80%) from 1,000 replicates are shown at internal nodes. Tip labels correspond to individual sample identifiers. The tree is drawn to scale, with branch lengths in the same units as those of the evolutionary distances to infer the phylogenomic tree. The evolutionary distances were computed using the Maximum Composite Likelihood Method and are in the units of the number of base substitutions per site. The analysis encompassed 168 sequences, aligned using MEGA12. Clades A–D were defined based on phylogenetic clustering. Colour indicates clade assignment as follows: blue (A), orange (B), green (C), and pink (D). The tree is rooted using the outgroup *Toxascaris leonina* MN329693 (scale bar: 0.02 nucleotide substitutions per base).

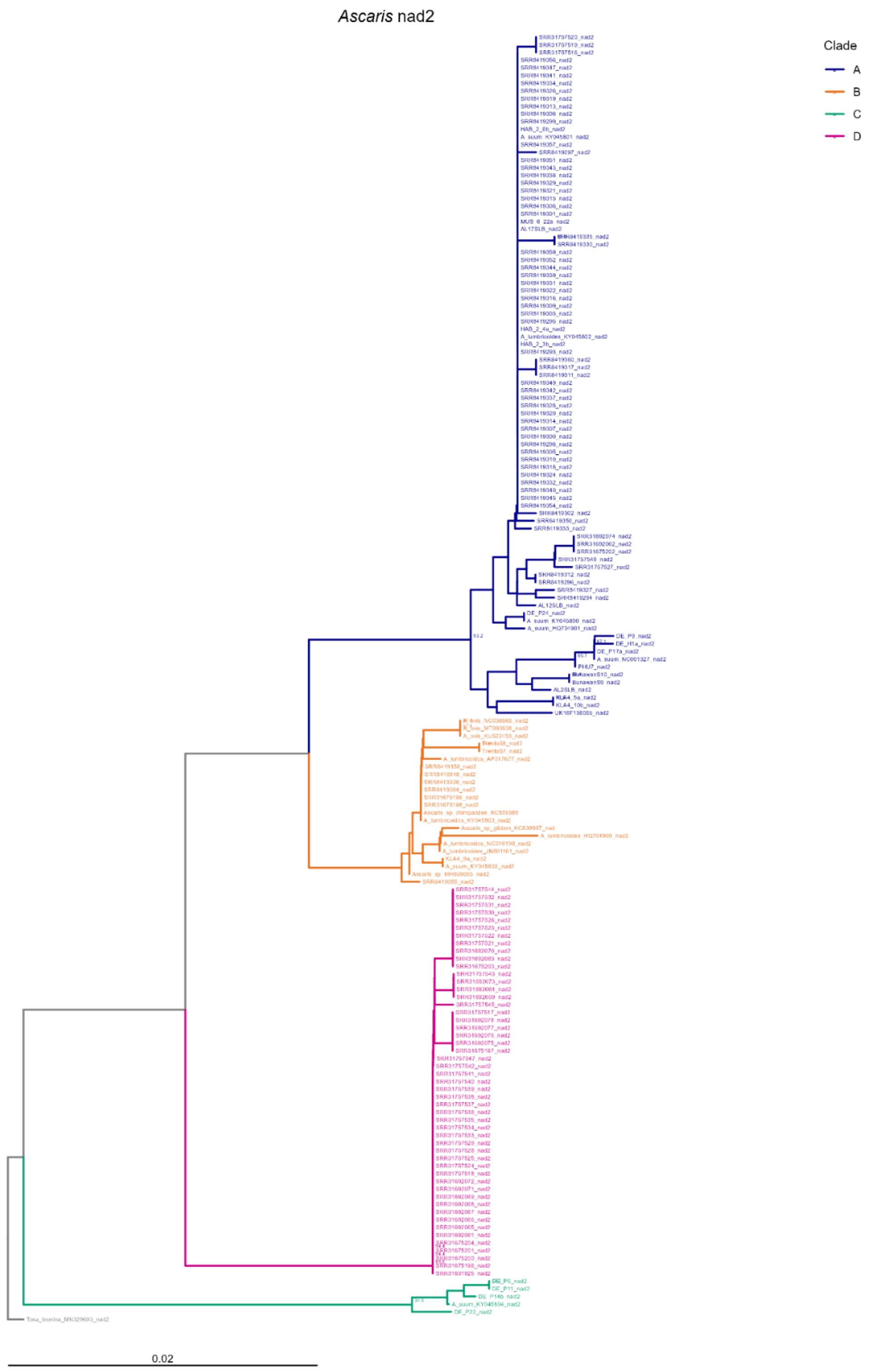


**Supplementary Figure S8** ***Neighbour-joining tree of nad2 from Ascaris samples.***
Bootstrap values (>80%) from 1,000 replicates are shown at internal nodes. Tip labels correspond to individual sample identifiers. The tree is drawn to scale, with branch lengths in the same units as those of the evolutionary distances to infer the phylogenomic tree. The evolutionary distances were computed using the Maximum Composite Likelihood Method and are in the units of the number of base substitutions per site. The analysis encompassed 168 sequences, aligned using MEGA12. Clades A–D were defined based on phylogenetic clustering. Colour indicates clade assignment as follows: blue (A), orange (B), green (C), and pink (D). The tree is rooted using the outgroup *Toxascaris leonina* MN329693 (scale bar: 0.02 nucleotide substitutions per base).

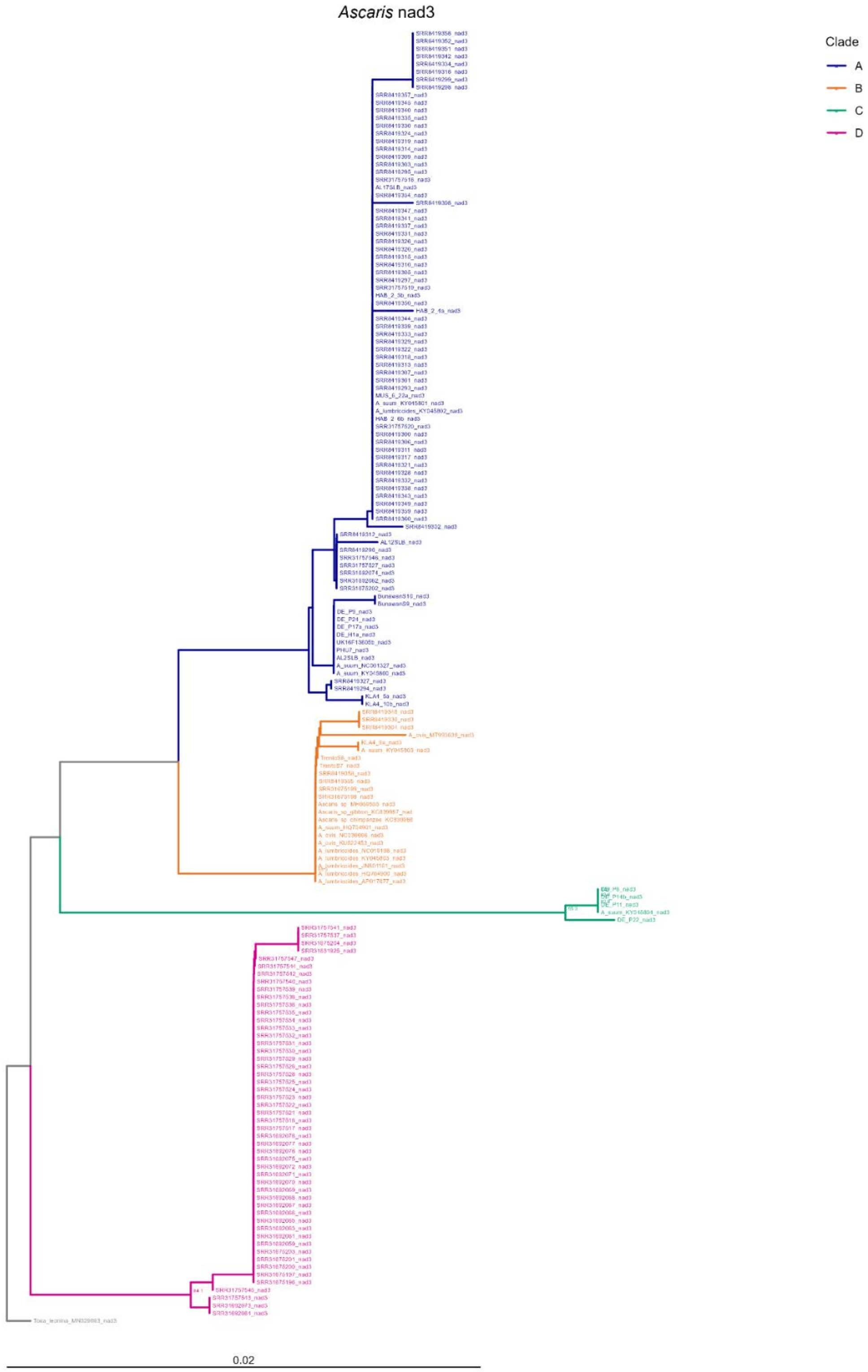


**Supplementary Figure S9** ***Neighbour-joining tree of nad3 from Ascaris samples.***
Bootstrap values (>80%) from 1,000 replicates are shown at internal nodes. Tip labels correspond to individual sample identifiers. The tree is drawn to scale, with branch lengths in the same units as those of the evolutionary distances to infer the phylogenomic tree. The evolutionary distances were computed using the Maximum Composite Likelihood Method and are in the units of the number of base substitutions per site. The analysis encompassed 168 sequences, aligned using MEGA12. Clades A–D were defined based on phylogenetic clustering. Colour indicates clade assignment as follows: blue (A), orange (B), green (C), and pink (D). The tree is rooted using the outgroup *Toxascaris leonina* MN329693 (scale bar: 0.02 nucleotide substitutions per base).

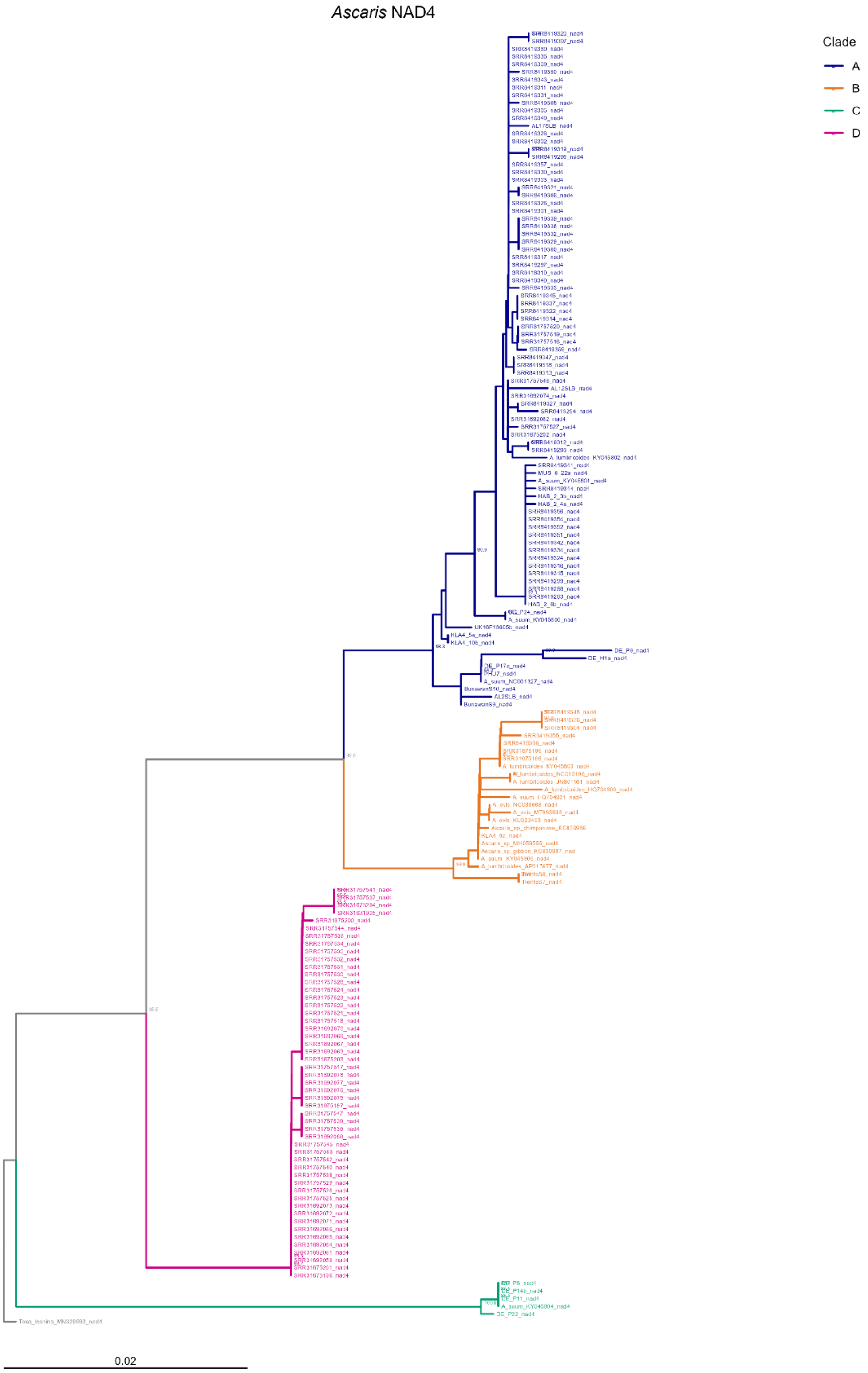


**Supplementary Figure S10** ***Neighbour-joining tree of nad4 from Ascaris samples.***
Bootstrap values (>80%) from 1,000 replicates are shown at internal nodes. Tip labels correspond to individual sample identifiers. The tree is drawn to scale, with branch lengths in the same units as those of the evolutionary distances to infer the phylogenomic tree. The evolutionary distances were computed using the Maximum Composite Likelihood Method and are in the units of the number of base substitutions per site. The analysis encompassed 168 sequences, aligned using MEGA12. Clades A–D were defined based on phylogenetic clustering. Colour indicates clade assignment as follows: blue (A), orange (B), green (C), and pink (D). The tree is rooted using the outgroup *Toxascaris leonina* MN329693 (scale bar: 0.02 nucleotide substitutions per base).

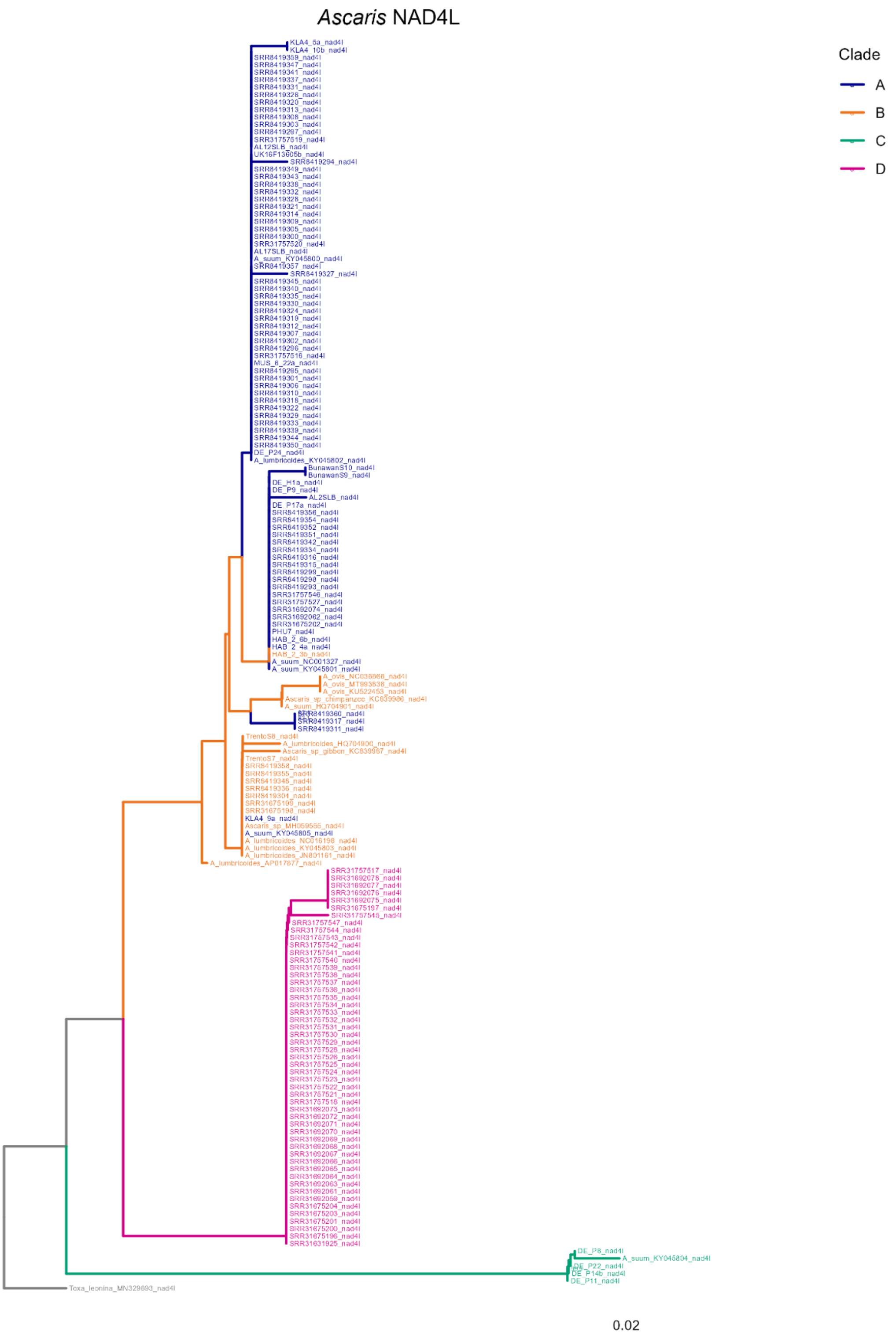


**Supplementary Figure S11** ***Neighbour-joining tree of nad4l from Ascaris samples.***
Bootstrap values (>80%) from 1,000 replicates are shown at internal nodes. Tip labels correspond to individual sample identifiers. The tree is drawn to scale, with branch lengths in the same units as those of the evolutionary distances to infer the phylogenomic tree. The evolutionary distances were computed using the Maximum Composite Likelihood Method and are in the units of the number of base substitutions per site. The analysis encompassed 168 sequences, aligned using MEGA12. Clades A–D were defined based on phylogenetic clustering. Colour indicates clade assignment based on concatenated dataset, which is as follows: blue (A), orange (B), green (C), and pink (D). The tree is rooted using the outgroup *Toxascaris leonina* MN329693 (scale bar: 0.02 nucleotide substitutions per base).

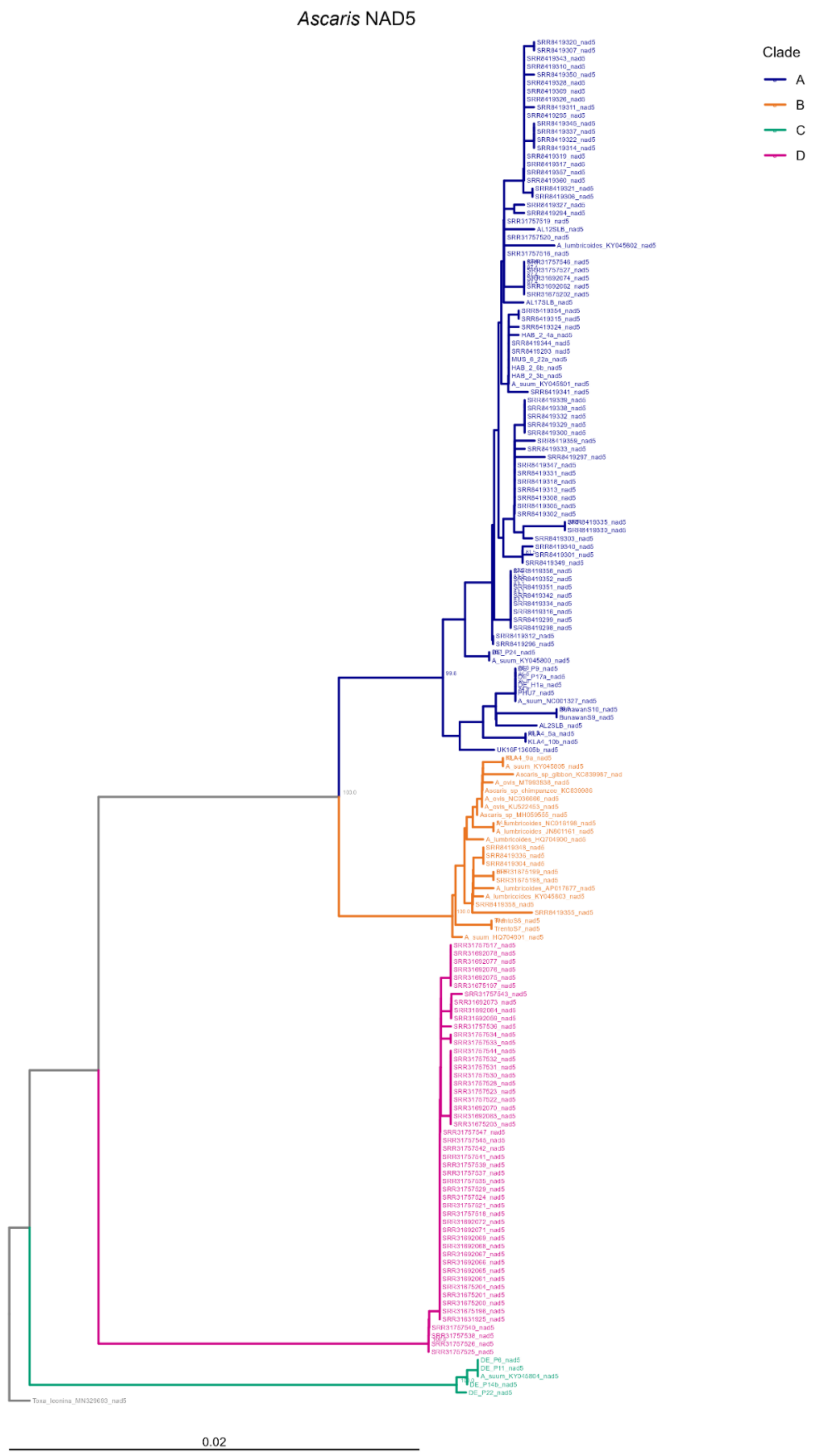


**Supplementary Figure S12** ***Neighbour-joining tree of nad5 from Ascaris samples*.**
Bootstrap values (>80%) from 1,000 replicates are shown at internal nodes. Tip labels correspond to individual sample identifiers. The tree is drawn to scale, with branch lengths in the same units as those of the evolutionary distances to infer the phylogenomic tree. The evolutionary distances were computed using the Maximum Composite Likelihood Method and are in the units of the number of base substitutions per site. The analysis encompassed 168 sequences, aligned using MEGA12. Clades A–D were defined based on phylogenetic clustering. Colour indicates clade assignment as follows: blue (A), orange (B), green (C), and pink (D). The tree is rooted using the outgroup *Toxascaris leonina* MN329693 (scale bar: 0.02 nucleotide substitutions per base).

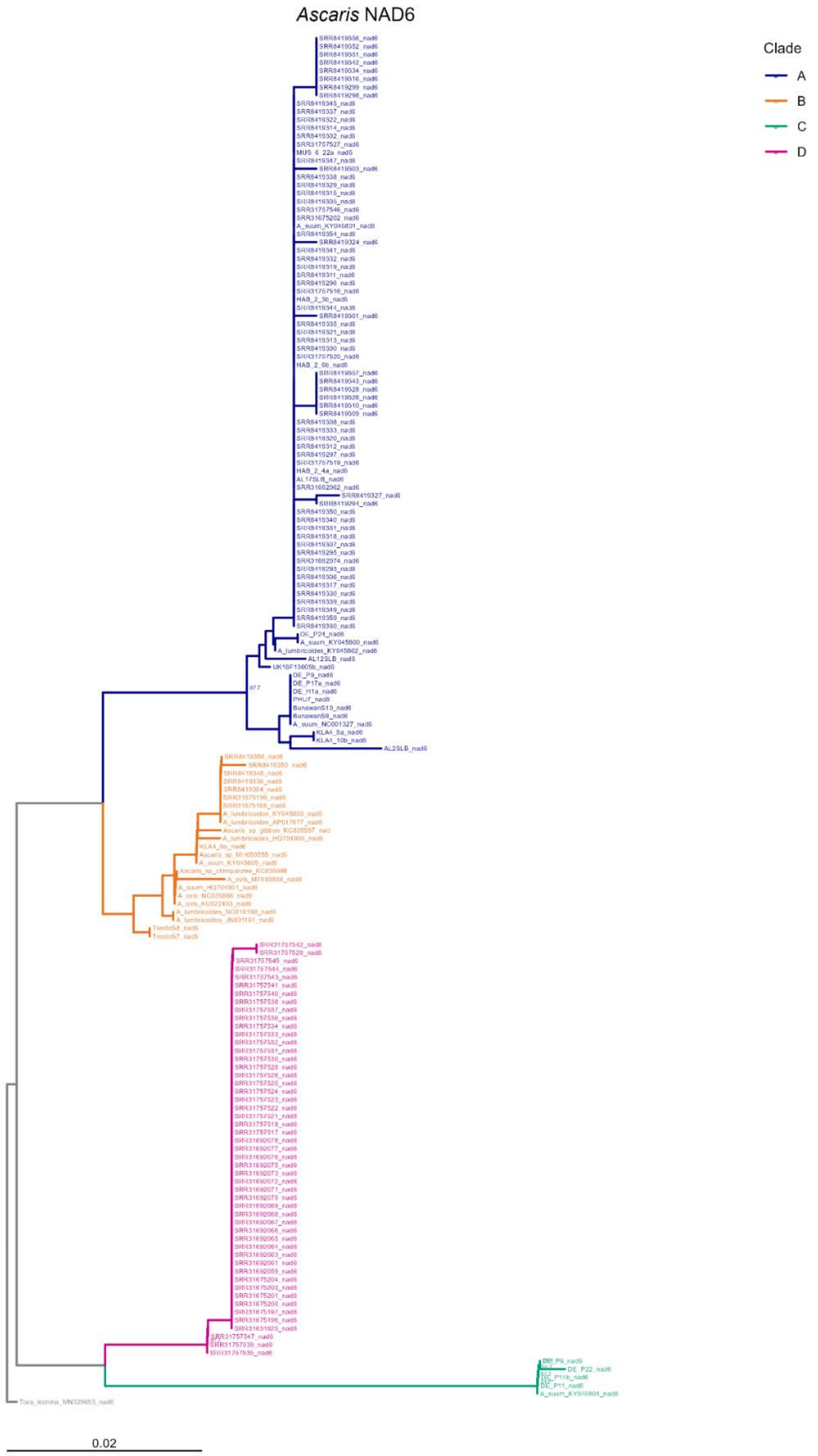


**Supplementary Figure S13** ***Neighbour-joining tree of nad6 from Ascaris samples.***
Bootstrap values (>80%) from 1,000 replicates are shown at internal nodes. Tip labels correspond to individual sample identifiers. The tree is drawn to scale, with branch lengths in the same units as those of the evolutionary distances to infer the phylogenomic tree. The evolutionary distances were computed using the Maximum Composite Likelihood Method and are in the units of the number of base substitutions per site. The analysis encompassed 168 sequences, aligned using MEGA12. Clades A–D were defined based on phylogenetic clustering. Colour indicates clade assignment as follows: blue (A), orange (B), green (C), and pink (D). The tree is rooted using the outgroup *Toxascaris leonina* MN329693 (scale bar: 0.02 nucleotide substitutions per base).
